# Hyper-Glycosylation as a Central Metabolic Driver of Alzheimer’s Disease

**DOI:** 10.1101/2025.04.30.651461

**Authors:** Tara R. Hawkinson, Zizhen Liu, Roberto A. Ribas, Terrymar Medina, Rikke S. Nielsen, Harrison A. Clarke, Xin Ma, Angela C. Mueller, Adrielle F. Plasencia, Alexander L. Shear, Samantha T. Simpson, Charles M. Soto, Jessica Sudderth, Feng Cai, Alex R. Cantrell, Matthieu G. Colpert, Cameron J. Shedlock, Lei Wu, Lyndsay E.A. Young, Damon D. Kooser, Li Chen, Alison M. Ryan, Roberto A. Ribas, Parastoo Azadi, Ralph J. Deberardinis, Stefan Prokop, Derek Allison, Yu Huang, Xing He, Jiang Bian, Craig W. Vander Kooi, Matthew S. Gentry, Ramon C. Sun

## Abstract

Alzheimer’s disease (AD) is a neurodegenerative disorder characterized by devastating degenerative decline. Metabolic disruptions are widely observed, yet their involvement in the molecular etiology of AD remains underexplored. Utilizing spatial metabolomics, lipidomics, and glycomics in both mouse models and human post-mortem samples, we identified a hyper-glycosylation phenotype as a hallmark of AD. To investigate the underlying mechanisms and whether the observed effect was a driver of the observed decline, we developed an advanced spatial isotopic tracing pulse-chase method to study the dynamics of N-linked glycans. Our analysis revealed enhanced glycan biosynthesis in AD mouse models. Based on these findings, we performed genetic and dietary interventions to modulate glycan biosynthesis. Genetic knockdown of glycan biosynthetic enzymes ameliorated the hyper-glycosylation and improved cognitive and behavioral outcomes in AD mice. In contrast, oral glucosamine supplementation drove hyper-glycosylation and exacerbated cognitive and behavioral deficits. To assess the clinical relevance of these findings, we conducted a retrospective analysis of a large population of patients with mild cognitive impairment (MCI), AD, and Alzheimer’s Disease Related Dementias (ADRD) stratified by glucosamine use, leveraging electronic health records. Consistently, glucosamine supplementation was associated with increased mortality in AD and ADRD patient cohorts, and significantly elevated progression from MCI to AD compared to age-matched controls. Collectively, our findings establish hyper-glycosylation as a pathological driver of AD and highlight glycan metabolism as an actional target in the fight against AD.

## Introduction

Alzheimer’s disease (AD) is a complex neurodegenerative disorder characterized by progressive cognitive impairment, synaptic dysfunction, neuroinflammation, and widespread neuronal loss^1,2^. Despite extensive research efforts elucidating underlying pathophysiology, effective disease-modifying treatments remain elusive^3^. Increasing evidence implicates metabolic dysfunction as a key component of AD pathology, with alterations in glucose metabolism^4^, mitochondrial function^5^, and lipid homeostasis^6^ playing contributory roles in disease progression. Fluorodeoxyglucose positron emission tomography (FDG-PET) imaging revealed significantly reduced glucose uptake in AD brains^7^. These FDG-PET changes are detectable years before clinical symptoms manifest, underscoring the importance of metabolic impairments in early disease pathology. Concurrently, lipid droplet accumulation has been identified as a metabolic hallmark of AD, suggesting that lipid dysregulation and energy homeostasis are tightly linked to neurodegeneration^8^. Recent studies demonstrating that restoring glucose metabolism can attenuate AD progression in animal models highlight the growing recognition of metabolic regulation as a critical frontier in AD research^9^. To this end, the entire metabolome and its changes have not been comprehensively explored in AD. For example, complex carbohydrate metabolism, such as glycan biosynthesis and processing, remains elusive in the context of AD. Understanding the overall metabolic shifts that occur in AD is essential for our understandings of AD disease progression.

The dynamic metabolome related to glucose utilization has not been defined in AD. The interconnected role of metabolites as not only sources of energy but as biosynthetic precursors in anabolic reactions remains unclear. In particular, complex carbohydrate metabolism including glycan biosynthesis and processing remains under defined in AD. N-linked glycosylation is a fundamental post-translational modification crucial for maintaining protein stability, intracellular trafficking, and receptor-ligand interactions critical for brain homeostasis ^10^. In the central nervous system, glycosylation plays a vital role in synaptic plasticity^11^, neurotransmitter receptor function^12^, and neuroimmune signaling^13^, making it integral to neuronal communication and resilience. Glycan metabolism is tightly linked to glucose^14^ and glucosamine availability^15^, forming a critical intersection between cellular energy balance and post-translational modifications. Connecting pathways, including the hexosamine biosynthetic pathway and glycan salvage pathways^16^, regulate glycosylation processes, further influencing protein folding, trafficking, and degradation. Aberrant glycan processing can lead to altered protein stability and dysfunctional cell signaling, contributing to neurodegenerative mechanisms^17^. Notably, congenital disorders of glycosylation (CDGs), caused by germline mutations in glycosylation-related genes, invariably result in neurological impairments, underscoring the essential role of glycosylation in brain development and function^18^. Alterations in glycosylation patterns have been documented across multiple neurological disorders^17^, suggesting a broader role for glycan modifications in neurodegeneration.

Recent AD research has revealed a complex interplay between multiple pathological processes, including the functional diversity of microglia subtypes^19^, tau propagation^20^, amyloid-beta and tau crosstalk^21^, neuroinflammation^22^, and lipid metabolism dysregulation^6^. Aberrant glycosylation could integrate or synergize with these pathological features, influencing multiple facets of AD progression. N-linked glycans regulate protein stability, intercellular communication, and immune signaling modification, is crucial for maintaining protein stability, intracellular trafficking, and receptor-ligand interactions^10^, and blood brain barrier regulation^23^, making them a crucial component of brain homeostasis. Disruptions in N-linked glycosylation can impact microglial function^24^, altering inflammatory responses and phagocytic capacity^25^. Additionally, glycan modifications influence tau post-translational modifications and aggregation dynamics^26^, potentially modulating tau propagation across neural networks. In the context of amyloid-beta pathology, glycosylation changes may alter plaque composition and clearance mechanisms^27^. Furthermore, glycan interactions with lipid metabolism contribute to lipid droplet dynamics^28^, influencing neuronal resilience to metabolic stress. Rather than acting as an isolated phenomenon, glycosylation represents an integrated metabolic event that intersects with multiple AD pathologies. Advances in spatial metabolomics, lipidomics, and glycomics provide powerful tools to dissect these intricate connections, offering new insights into the role of glycan dysregulation in AD pathogenesis.

Building on these advancements, we hypothesized that hyper-glycosylation is a pathological hallmark of AD that actively contributes to disease progression. While prior studies have suggested glycosylation changes in AD^29^, the functional implications of altered glycan biosynthesis remain unclear. To address this gap, we employed an integrative approach combining spatial metabolomics^30,31^ and advanced spatial isotopic tracing^32,33^ to elucidate glycan dynamics in AD mouse models and human post-mortem brain tissue. Using pulse-chase labeling of complex glycans, we uncovered increased glycan biosynthesis in AD brains, indicative of a metabolic shift toward hyper-glycosylation. Given the extensive involvement of glycans in maintaining brain function, these findings raise critical questions regarding the influence of glycosylation on AD homeostasis and disease pathophysiology. To investigate the functional impact of hyper-glycosylation, we conducted both genetic and dietary interventions aimed at modulating glycan biosynthesis. Genetic knockdown of key glycosylation enzymes ameliorated cognitive deficits in AD mice, while oral glucosamine supplementation exacerbated behavioral impairments, supporting a causal role for glycan dysregulation in disease severity. These findings suggest that hyper-glycosylation is not simply a secondary feature of neurodegeneration but may function as a critical driver of AD pathology. To extend these insights to human populations, we performed a retrospective analysis of mild cognitive impairment (MCI) and dementia patients stratified by glucosamine supplementation using electronic health records^34^. Consistently, glucosamine use was associated with increased mortality and a higher likelihood of progression from MCI to AD, emphasizing the potential risks of excessive glycan biosynthesis in disease advancement. In summary, our findings establish hyper-glycosylation as a significant driver of AD progression and establish the potential of targeting glycan biosynthesis as a novel therapeutic strategy.

## Results

### Hyper-glycosylation in human AD specimens

Our laboratory developed and implemented a multiplexed matrix-assisted laser desorption ionization (MALDI)-mass spectrometry imaging (MSI) approach to conduct spatial metabolomics, lipidomics, and glycomics on a single mouse brain tissue section^31^. Initially, we applied this methodology to human Alzheimer’s disease (AD) brain samples. However, when comparing normal and AD human brain tissues, we observed markedly low glycan signal intensity with poor spatial resolution (supplemental Fig. 1). Notably, lipid content remained elevated in human samples even following Carnoy’s solution treatment (ethanol, acetic acid, and chloroform in a 60:30:10% ratio), a protocol conventionally sufficient for lipid removal in mouse brain tissue and widely utilized in MALDI-based analyses^35^. To enhance spatial N-glycome detection, we investigated alternative lipid extraction methodologies to improve glycan signal recovery in human brain specimens. We implemented an optimized xylene wash protocol (one-hour washes, repeated three times), which proved to be the effective lipid removal method and significantly enhanced glycan signal intensity (supplemental Fig. 1A).

Sequential MALDI imaging across multiomics datasets confirmed that this approach facilitated robust spatial glycomics analysis, allowing for a more comprehensive assessment of glycan distribution with over 10-20-fold increase in glycan intensities in AD and control tissues (supplemental Fig. 1B-C). Using this optimized multiomics workflow, we performed spatial metabolomics, lipidomics, and glycomics in a cohort of human frontal cortex samples from normal and AD patients (n=3 each), matched for age, sex, and post-mortem interval (Fig. 1A). Metabolomics and lipidomics analyses revealed specific AD-associated molecular alterations, including both upregulated and downregulated species (supplemental Fig. 2-3). However, spatial glycomics analysis uncovered a particularly striking pattern: glycan abundance was dramatically elevated across both white and gray matter regions in AD samples (Fig. 1B and Extended Fig. 1). For example, both the bisecting glycans (1688 m/z) and high mannose (1419 m/z) are elevated in the grey matter region of the AD brain (Fig. 1C). Further spatial metabolomics analysis revealed a significant reduction in N-acetylglucosamine and glucosamine-6-phosphate, both critical precursors in N-glycan biosynthesis (Fig. 1D). The unexpected and widespread increase in glycosylation underscored the need for mechanistic studies to elucidate the biochemical pathways driving this altered glycan profile and its implications in AD pathology.

**Fig. 1.**
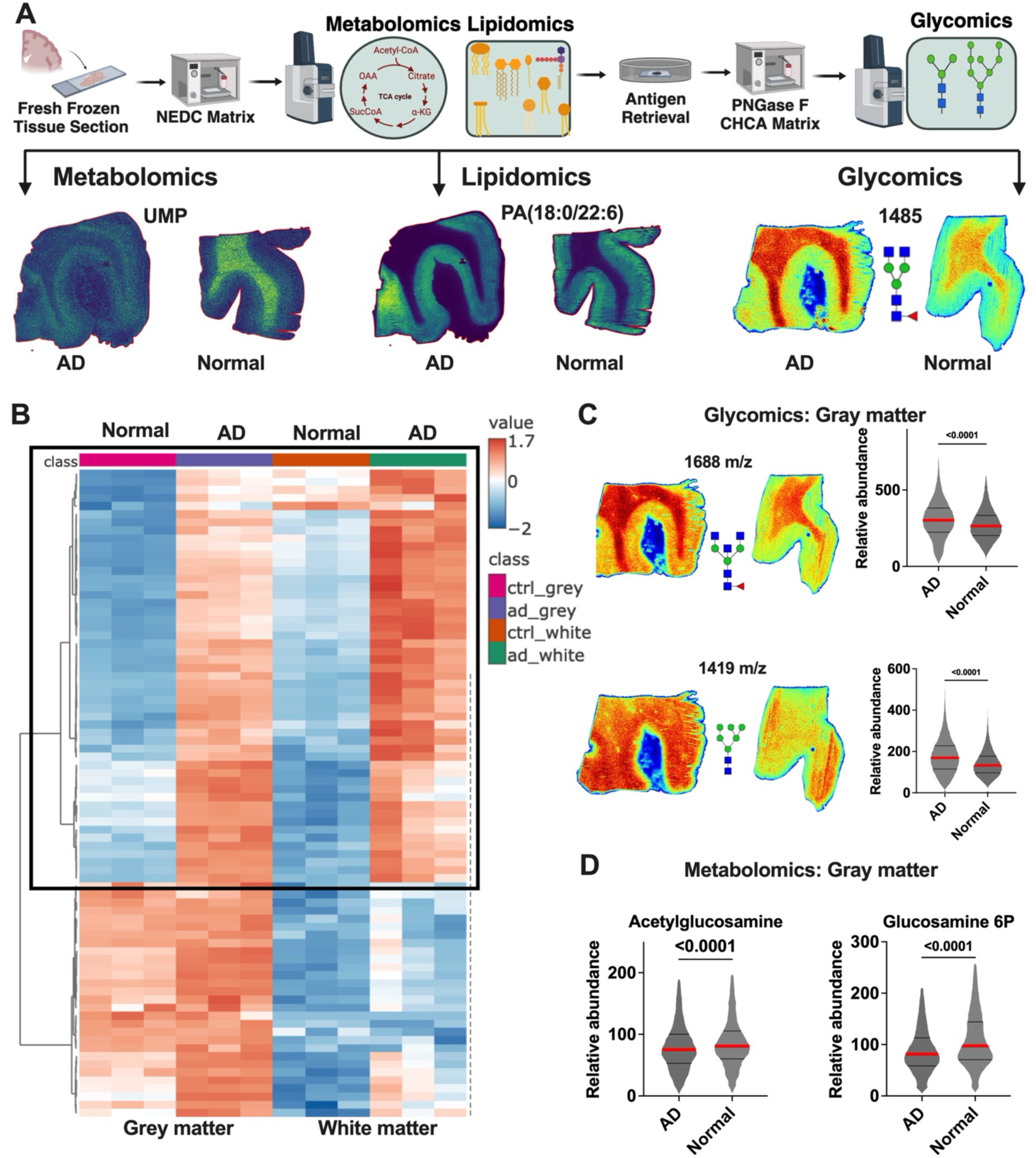
Spatial multiomics analysis reveals hyper-glycosylation in human Alzheimer’s disease brains. **a**, Schematic overview of the multiplexed mass spectrometry imaging (MSI) workflow utilized for integrated spatial metabolomics, lipidomics, and glycomics from human brain tissue sections. Representative images for metabolite uridine monophosphate (UMP), lipid (PA(18:0/22:6)), and glycan (m/z 1485) distributions in Alzheimer’s disease (AD) and normal control brains are shown. **b**, Heatmap illustrating the differential abundance of glycan species identified by spatial glycomics analysis, comparing grey and white matter regions from control (Normal) and AD brain specimens (n=3, each)..**c**, Spatial glycomics visualization and quantification of representative N-glycan species (1688 m/z and 1419 m/z) in grey matter regions. Violin plots show significantly increased abundance of these glycans in AD relative to normal controls. Statistical analysis was performed using two-tailed t-tests (n>2000 pixels per group); exact P-values indicated. **d**, Spatial metabolomics analysis in grey matter regions reveals a significant reduction in key glycan precursors, acetylglucosamine and glucosamine-6-phosphate, in AD compared to normal control samples. Violin plots present relative abundances; statistical significance assessed by two-tailed t-tests (n>2000 pixels per group); exact P-values indicated.

To investigate the mechanistic basis of hyper-glycosylation in Alzheimer’s disease (AD), we first sought to validate this phenotype in a broader cohort of human AD specimens. We assembled an additional set of AD patient samples, ensuring matched controls for age and post-mortem interval (PMI) while stratifying individuals by Braak staging (Fig. 2A-B). This stratification allowed assessment of whether hyper-glycosylation follows a temporally regulated trajectory throughout disease progression. We utilized formalin-fixed, paraffin-embedded (FFPE) patient samples (Fig. 2A). Spatial glycomics workflows incorporating PNGase-mediated glycan release and MALDI-based mass spectrometry imaging enabled the quantification of regional glycan distribution matched to histological annotation (Fig. 2A and B). Analysis of glycosylation in both grey and white matter revealed distinct patterns of stage-dependent alterations. In grey matter, a heatmap of glycan abundance demonstrated a progressive increase in N-linked glycosylation across Braak stages, with the most pronounced changes observed in later stages (Figure 2C). Violin plots of glycan structures at 1688 m/z and 1409 m/z further illustrated these trends, with a steady increase in glycosylation across Braak stages (Figure 2D). Spatial heatmaps confirmed these findings, showing localized glycan accumulation patterns that intensified with AD pathology (Figure 2E). White matter glycosylation exhibited a different pattern, with glycan abundance increasing at Braak stages 1-2 but not persisting through later stages (Extended Figure 2A). A heatmap of glycan abundance highlighted this transient elevation, suggesting that hyper-glycosylation in white matter may be an early but non-progressive event. Violin plots of glycan structures at 1079 m/z and 1905 m/z indicated an early stage rise in white matter glycosylation without sustained accumulation at later stages (Extended Figure 2B and C). These findings suggest that hyper-glycosylation in grey matter is more progressive and associated with AD severity. These results validate the reproducibility of the hyper-glycosylation phenotype and implicating N-glycan modifications as a potential driver of AD pathology. The next critical step is to delineate the metabolic mechanisms underlying this phenotype.

**Fig. 2.**
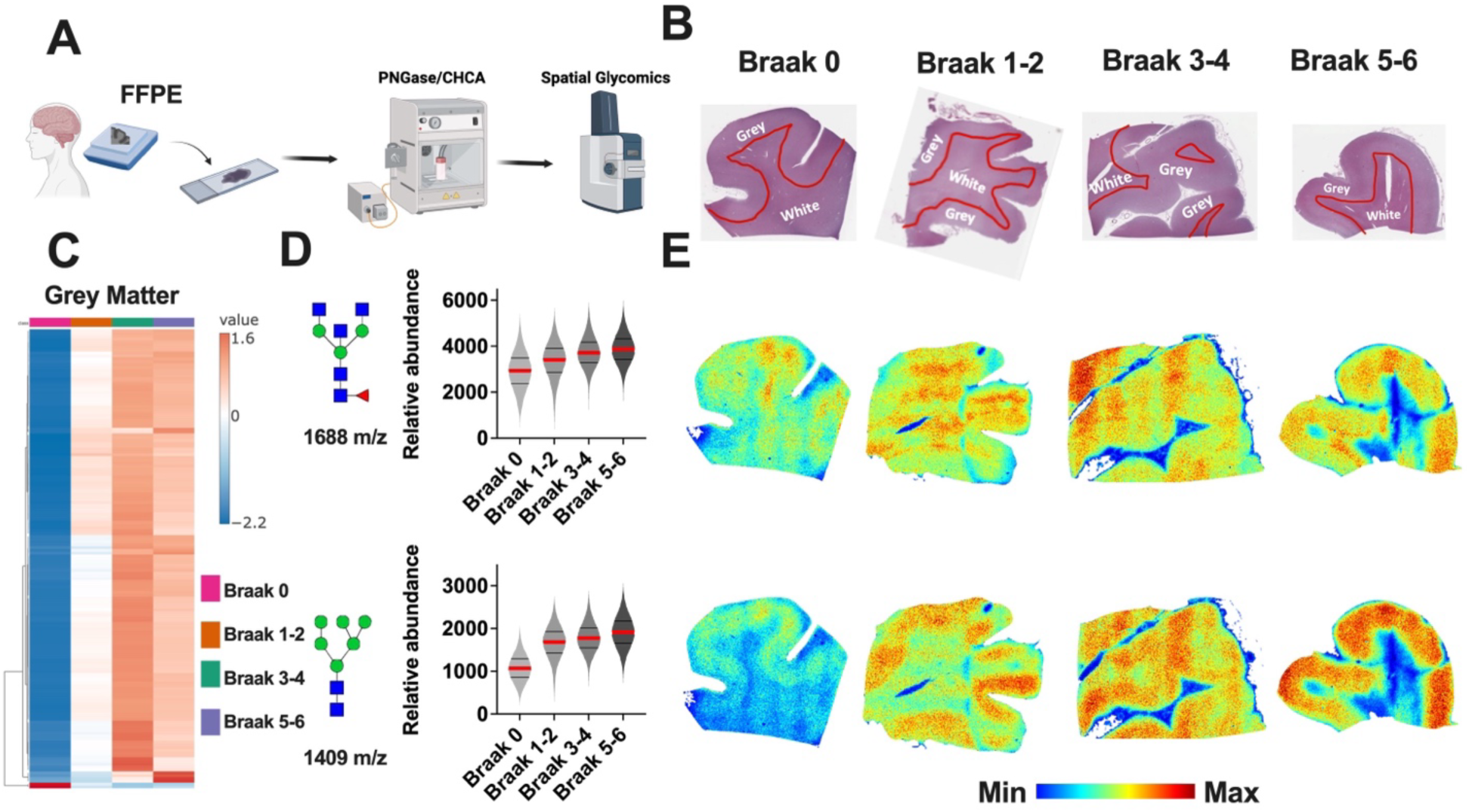
Progressive hyper-glycosylation across Braak stages in human Alzheimer’s disease. **a**, Schematic illustrating the workflow for spatial glycomics using formalin-fixed, paraffin-embedded (FFPE) human brain samples, incorporating PNGase treatment followed by CHCA matrix application for MALDI imaging. **b**, Representative histological sections indicating anatomical delineation of grey and white matter regions from human brain tissues stratified by Braak stages (0, 1-2, 3-4, and 5-6). **c**, Heatmap displaying hierarchical clustering of glycan abundance specifically in grey matter regions across Braak stages, illustrating progressive glycan accumulation correlated with advancing disease severity. **d**, Violin plots representing quantitative glycan analysis of two specific glycans (1688 m/z and 1409 m/z) in grey matter across Braak stages. Data indicate a progressive increase in glycan abundance with disease advancement. **e**, Spatial glycomics images demonstrating the distribution and intensity of glycans (1688 m/z and 1409 m/z) in human brain samples across Braak stages. Spatial intensity increases correlate with disease severity, prominently highlighting regional glycan accumulation.

**Fig. 3.**
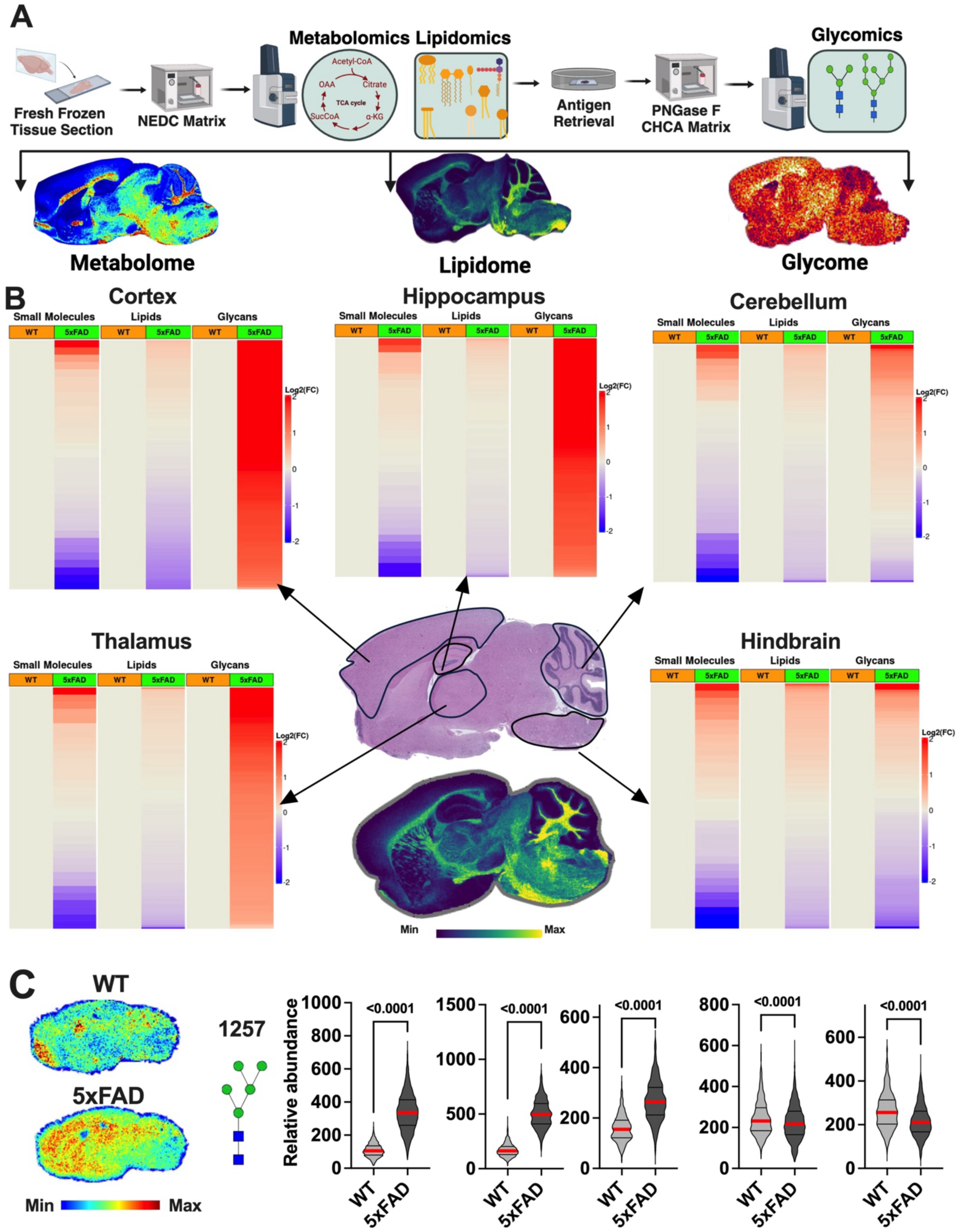
Conserved hyper-glycosylation in the 5xFAD mouse model of Alzheimer’s disease. **a**, Schematic representation of the multiomics workflow employed for spatial metabolomics, lipidomics, and glycomics analysis on fresh frozen brain tissue sections from wild-type (WT) and 5xFAD mice, illustrating representative metabolite, lipid, and glycan spatial distributions. **b**, Comparative heatmaps of metabolite, lipid, and glycan abundances in distinct brain regions (cortex, hippocampus, cerebellum, thalamus, and hindbrain) from WT and 5xFAD mice. WT levels are set to 1 while Log2 fold changes indicate significant alterations across regions, with pronounced increases in glycan abundance specifically observed in 5xFAD brains in the cortex, hippocampus and thalamus (n>2000 pixels per group). **c**, Spatial glycomics visualization of a representative high mannose glycan (m/z 1257) distribution, demonstrating marked elevation in the 5xFAD model compared to WT. Corresponding violin plots quantify glycan abundance across multiple brain regions, confirming statistically significant increases in 5xFAD mice. Statistical analysis performed using two-tailed t-tests; exact P-values indicated.

### Hyper-glycosylation in mouse AD brains

To investigate the metabolic mechanisms underlying hyper-glycosylation, we sought to determine whether the 5xFAD mouse model of AD replicates the hyper-glycosylation phenotype observed in human AD brains. To address this, we performed spatial metabolomics, lipidomics, and glycomics in 9-month-old wild-type (WT) and 5xFAD mouse brains (Fig. 3A), corresponding to a stage of significant disease manifestation^36^.

Consistent with human AD samples, we identified widespread alterations in the metabolome and lipidome of 5xFAD mice (Fig. 3B and Extended Fig. 3A), corroborating previous studies that have linked AD pathology to disrupted energy metabolism and lipid homeostasis^37,38^. Notably, N-glycan profiling revealed the same hyper-glycosylation phenotype in the 5xFAD mouse model as observed in human AD specimens (Fig. 3B and Extended Fig. 3D). Further global spatial analysis demonstrated that hyper-glycosylation was regionally specific, being most pronounced in the cortex, hippocampus, and thalamus, while being less prominent in the cerebellum and hindbrain (Fig. 3C and Extended Fig. B-C). These findings suggest that hyper-glycosylation preferentially affects regions associated with memory, cognitive processing, and neuroinflammation, aligning with known patterns of neurodegeneration in AD. Taken together, these results indicate that hyper-glycosylation is a conserved feature of AD pathology across species and suggest that specific brain regions may be differentially affected by this metabolic dysregulation.

### Increased glycan biosynthesis in mouse model of AD

Having established that hyper-glycosylation is a conserved phenotype in both human AD brains and major regions of the 5xFAD mouse model, it is critical to elucidate the underlying metabolic mechanisms driving hyper-glycosylation. Hyper-glycosylation can arise from either increased glycan biosynthesis within the endoplasmic reticulum (ER) and Golgi apparatus^39^ or reduced degradation and recycling through the lysosomal pathway^16^. To distinguish between these possibilities, we employed a stable isotope tracing approach coupled with a pulse-chase experiment^40^. The pulse phase provides a direct measure of *de novo* glycan biosynthesis, whereas the chase phase enables an assessment of glycan turnover and salvage activity (Fig. 4A). To facilitate deep metabolic tracing, we implemented a ^13^C-enriched liquid diet^41^, an approach that allows extensive isotopic labeling of glycans given their terminal position in metabolic pathways. Unlike small metabolic intermediates, which incorporate relatively few labeled carbons, N-glycans can integrate up to 40–60 labeled carbon atoms derived from glucose, posing challenges in resolving individual isotopologues. To address this challenge, we leveraged the ion mobility capabilities of our MALDI system^33,42^, which discriminates molecular species based on their shape and size in addition to mass. This approach enables accurate identification of ^13^C-labeled glycans, as isotopologues exhibit predictable mass shifts while maintaining identical collision cross-section (CCS) values, facilitating precise molecular characterization (Fig. 4B).

**Fig. 4.**
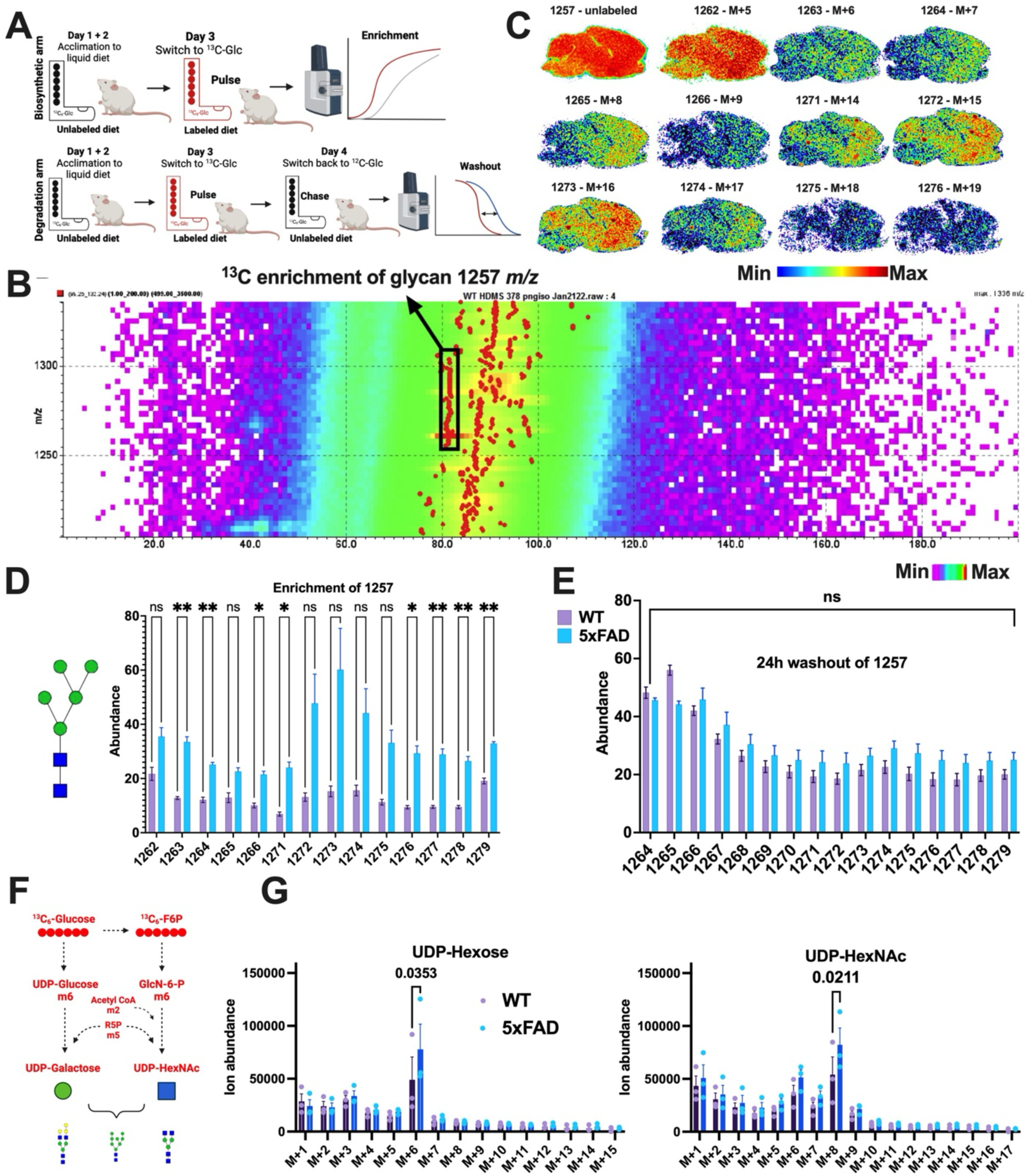
Increased glycan biosynthesis drives hyper-glycosylation in the 5xFAD mouse model. **a**, Schematic of the stable isotope tracing experimental design used to differentiate glycan biosynthesis (pulse phase) from glycan degradation and recycling (chase phase) in wild-type littermates (WT) and 5xFAD mice, employing a 13C6-glucose enriched liquid diet. **b**, Ion mobility mass spectrometry imaging (MALDI) showing 13C6 enrichment of the glycan species at 1257 m/z, highlighting distinct isotopologue separation. **c**, Spatial distributions of unlabeled and 13C6-labeled isotopologues of the glycan (m/z 1257) across the mouse brain tissue section, demonstrating progressive labeling. **d**, Quantitative analysis comparing the enrichment levels of various isotopologues of glycan 1257 m/z in WT and 5xFAD brains during the pulse phase, indicating significantly enhanced glycan biosynthesis in 5xFAD mice. Statistical significance determined by two-tailed t-tests; ns, not significant; *P<0.05, **P<0.01. **e**, Analysis of glycan isotopologue abundances following a 24-hour washout period (chase phase), revealing no significant differences in glycan turnover between WT and 5xFAD mice. **f**, Metabolic schematic illustrating the incorporation pathway of 13C6-glucose into UDP-Hexose and UDP-HexNAc, key glycan biosynthetic precursors. **g**, LC-MS-based isotope tracing analysis confirming increased levels of labeled UDP-Hexose (left) and UDP-HexNAc (right) in 5xFAD mice relative to WT controls (N=3 individual samples each genotype), supporting elevated glycan biosynthesis. Statistical significance assessed by two-tailed t-tests; exact P-values indicated.

For the experimental, one cohort of mice was administered a ^13^C-labeled liquid diet for 24 hours to measure biosynthetic flux (pulse phase), while a second cohort received the same diet followed by a 48-hour washout period to evaluate glycan turnover (chase phase) (Fig. 4A). Spatially resolved MALDI imaging revealed distinct distributions of both native (M0) and labeled glycan species, confirming successful isotopic incorporation and metabolic tracing (Fig. 4B and C). Comparative analysis between WT and 5xFAD mice demonstrated a significant increase in glycan enrichment in 5xFAD mice, particularly in high mannose glycans such as the species with m/z 1257 (Fig. 4D). This enrichment was evident across multiple isotopologues in 5xFAD mice relative to WT controls (Extended Fig. 4A-C), whereas no significant differences were observed following the 48-hour washout period (Fig. 4E and Extended Fig. 4A-C). These data revealed that increased glycan biosynthesis, rather than impaired glycan degradation or salvage, is the predominant driver of hyper-glycosylation in AD pathology.

To further substantiate these findings, we performed pooled LC-MS-based isotope tracing from the same experimental cohorts. This analysis revealed a marked increase in labeled UDP-hexose and UDP-GlcNAc (Fig. 4F-G), two key precursors in the N-glycan biosynthetic pathway^43^. To further confirm that glycan biosynthesis is increased in AD, we analyzed the expression of glycan biosynthetic enzymes in both human and mouse AD models (Extended Fig. 5 and supplemental Fig. 4). Using RT-PCR and digital PCR, we quantified mRNA levels of key glycan biosynthesis genes in the ER and Golgi, including Mgat, Man1a2, and B4galt1. In both human AD brains and 5xFAD mouse models, we observed upregulation of these genes compared to their respective controls (Extended Fig. 5 and supplemental Fig. 4). These findings further support the notion that increased glycan biosynthesis is a key factor driving hyper-glycosylation in AD.

**Fig. 5.**
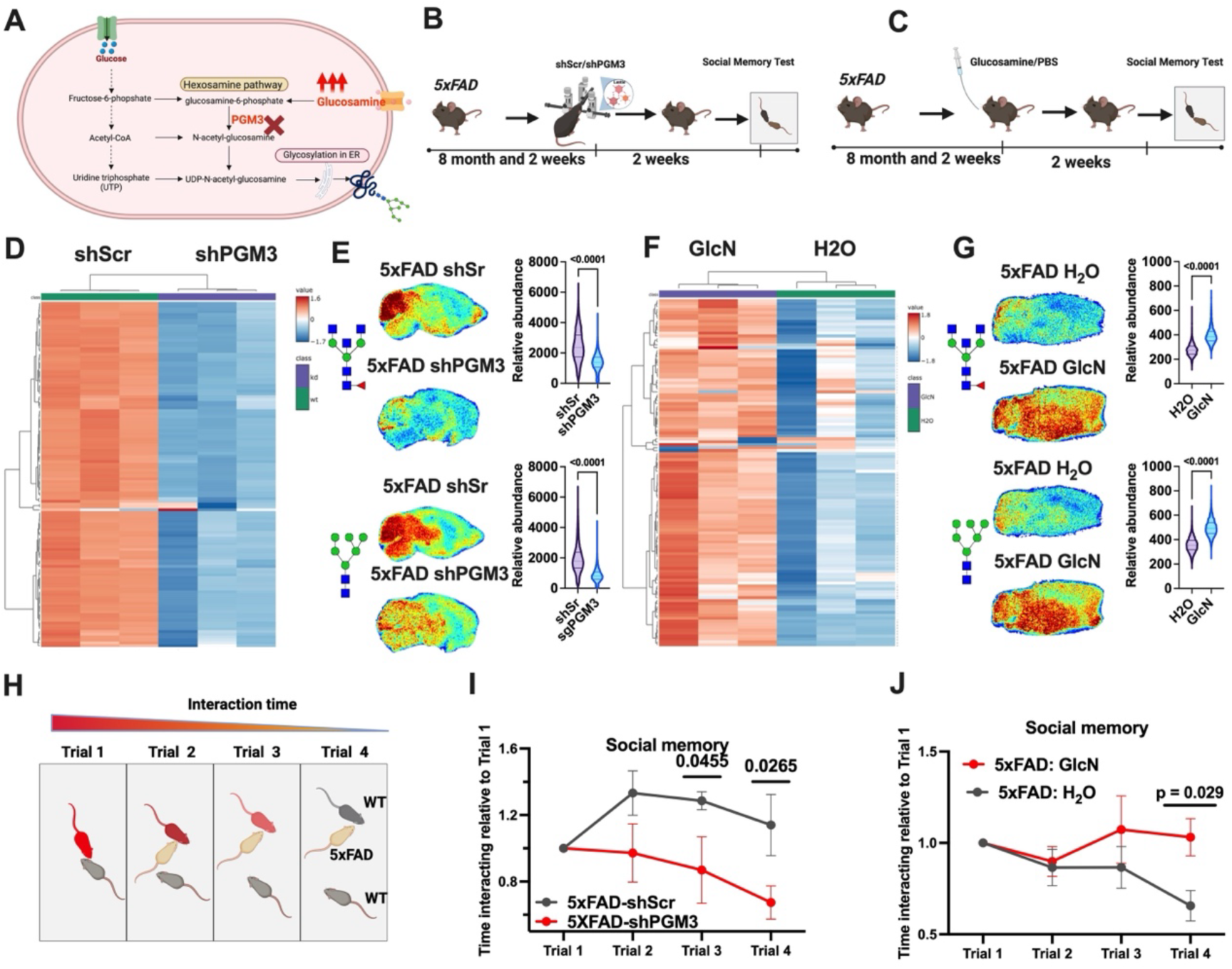
Genetic and dietary modulation of glycan biosynthesis impacts cognitive function in 5xFAD mice. **a**, Diagram illustrating the hexosamine biosynthetic pathway highlighting the intervention point at phosphoglucomutase 3 (PGM3) and glucosamine to modulate glycosylation. **b-c**, Experimental design schemes showing the genetic knockdown of PGM3 via shRNA delivery (b) and dietary modulation using oral glucosamine (GlcN) supplementation (c), both followed by a social memory test. **d**, Heatmap demonstrating global glycan abundance reduction in brains of 5xFAD mice treated with shPGM3 via lentivirus compared to scrambled control (shScr). **e**, Spatial glycomics images and corresponding violin plots quantifying significant reductions in glycan abundance in shPGM3-treated mice relative to controls. Statistical significance assessed by two-tailed t-tests. **f**, Heatmap depicting global glycan abundance increases in glucosamine (GlcN)-supplemented 5xFAD mice compared to water gavaged (H_2_O) controls. **g**, Spatial glycomics visualization and quantification indicating significant elevations in glycan abundance in GlcN-treated mice versus controls. Statistical significance assessed by two-tailed t-tests. (N=3 mice per group) **h**, Illustration of the social memory test paradigm, with interaction time expected to decrease over repeated exposures to familiar conspecifics. **i-j**, Social memory test results showing cognitive improvement in shPGM3-treated mice (i) and exacerbated cognitive deficits in glucosamine-supplemented mice (j). Data analyzed using repeated measures ANOVA; exact P-values indicated.

### Glycoproteomic Analysis

The observed hyper-glycosylation phenotype raised the question of whether this increase is due to the emergence of novel glycosylated proteins or an upregulation of glycosylation modifications on pre-existing proteins. To address this question, we conducted glycoproteomic analysis in both human and mouse samples^44^. Similar to our glycan enzyme transcriptomics approach, we analyzed pooled samples of human normal and AD frontal cortical tissues as well as WT and 5xFAD mouse brains. Given the necessity of enriching membrane glycoproteins prior to mass spectrometry analysis, we pooled n=4 samples per group to ensure sufficient input material. Membrane proteins were first extracted and enriched using high-salt and carbonate-based fractionation, followed by sequential ultracentrifugation.

Isolated membrane fractions were denatured, reduced, alkylated, and subjected to protease digestion. Glycopeptides were subsequently analyzed using nano-LC coupled to a high-resolution Thermo Eclipse mass spectrometer (Extended Fig. 6 and supplemental Fig. 5). Glycoproteomic datasets were processed using Proteome Discoverer (v3.0), with glycoproteins identified through Byonic database searches.

**Fig. 6.**
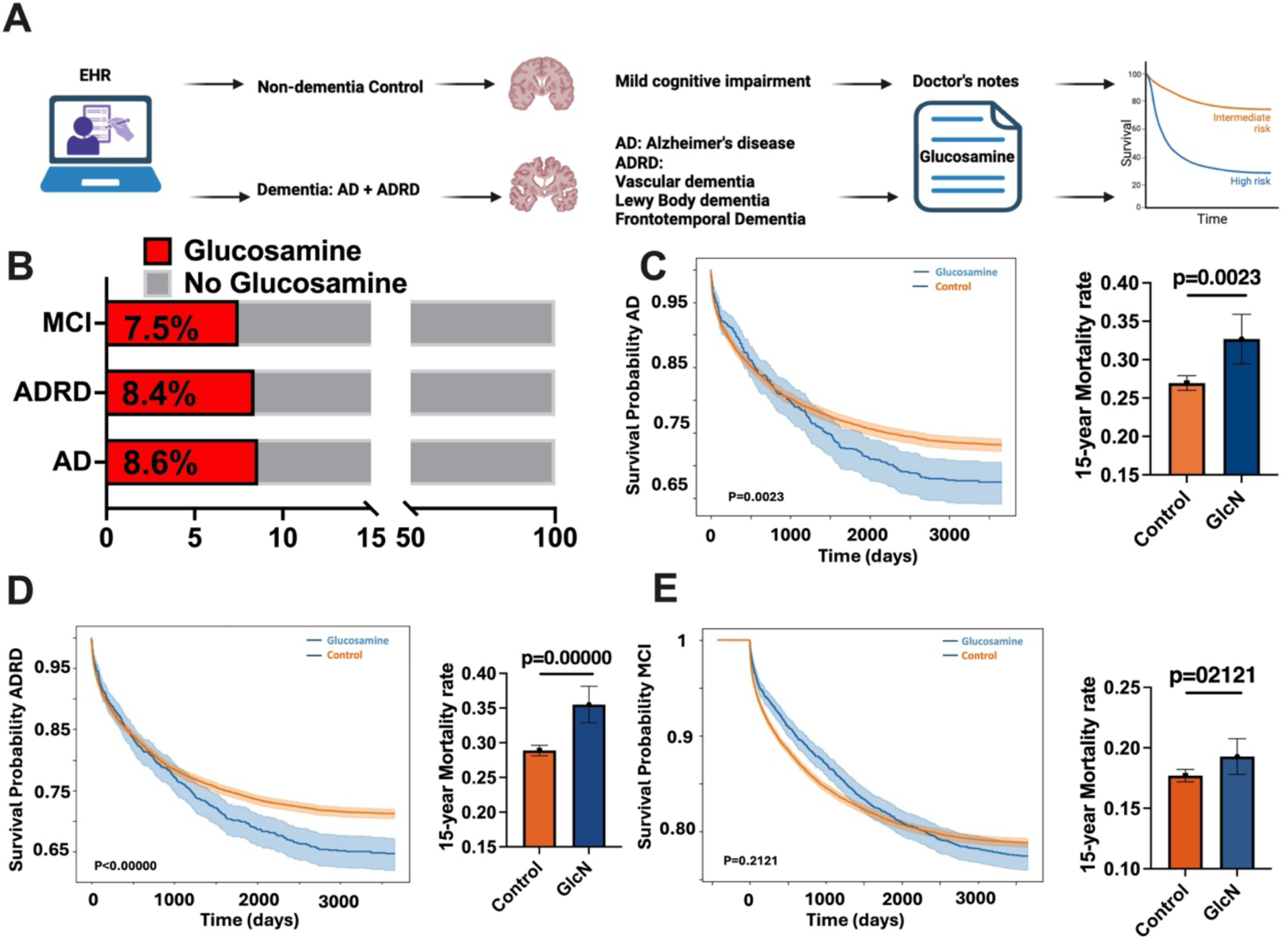
Real-world impact of glucosamine supplementation on Alzheimer’s disease patient outcomes. **a**, Schematic illustration of electronic health record (EHR)-based retrospective analysis design, categorizing patient cohorts into Alzheimer’s disease (AD), Alzheimer’s disease-related dementias (ADRD), and mild cognitive impairment (MCI) for survival analysis. **b**, Bar graph depicting the proportion of patients within MCI, ADRD, and AD cohorts identified as glucosamine users based on documented medical records. **c-e**, Kaplan-Meier survival curves and associated 15-year mortality rates comparing glucosamine users and non-users within AD (c), ADRD (d), and MCI (e) patient populations. Statistical analysis using log-rank test; exact P-values indicated.

From mouse samples, we identified 137 unique glycopeptides (Extended Fig. 6), while human extracts yielded 77 unique glycopeptides (supplemental Fig. 5). Relative quantification revealed a significant increase in glycopeptide abundance in AD samples compared to their respective WT controls (Extended Fig. 6B and supplemental Fig. 5B), further supporting the hyper-glycosylation phenotype identified via MALDI. Notably, 55% and 75% of glycopeptides were shared between AD and normal in human and mouse samples respectively (Extended Fig. 6C and supplemental Fig. 5C), respectively. Since multiple glycopeptides can be derived from the same protein, we observed over 90-95% overlap in glycoprotein identities between normal and AD samples (supplemental data 1). These findings indicate that hyper-glycosylation in AD predominantly involves an increase in glycosylation modifications on pre-existing glycoproteins rather than the appearance of novel glycosylated proteins. Finally, we conducted cell-type enrichment analysis through WebCSEA^45^, based on the identified glycopeptides to infer which neuronal populations are most affected by hyper-glycosylation. Our analysis suggests that neurons are the predominant cell population undergoing enhanced glycosylation, underscoring the potential role of aberrant glycosylation in neuronal dysfunction and AD pathogenesis (Extended Fig. 6D-E and supplemental Fig. 5D-E).

### Genetic and Dietary Interventions of Glycosylation During Neurodegeneration

To determine whether hyper-glycosylation is a causal driver of neurodegeneration or a secondary consequence of disease pathology, we implemented both genetic and dietary strategies to selectively decrease and increase N-glycan levels in the brain. This approach allowed us to assess the functional impact of altered glycosylation on neurobehavioral outcomes and neuropathological markers in AD (Fig. 5A). To suppress glycosylation, we targeted phosphoglucomutase 3 (PGM3), a key enzyme in the hexosamine biosynthetic pathway responsible for converting glucosamine-6-phosphate to N-acetylglucosamine-6-phosphate^46^. By selectively inhibiting PGM3, we aimed to attenuate glycan biosynthesis while minimizing off-target metabolic perturbations in the brain since targeting upstream enzymes in the pathway would disrupt central glucose metabolism, potentially leading to severe neurotoxic effects or lethality^47^. In complement, to enhance glycosylation levels, we administered glucosamine via oral gavage. Glucosamine is known to readily cross the blood-brain barrier and incorporate directly into brain glycans^48–50^, making it an ideal substrate for investigating the effects of increased glycosylation. If hyper-glycosylation is an active contributor to AD pathology, we hypothesized that glucosamine supplementation would exacerbate neurodegenerative phenotypes, whereas PGM3 inhibition would have protective effects.

To test the effects of glycosylation modulation, 8-month-old female 5xFAD mice received either shScrambled or shPGM3 packaged in lentivirus^51^ delivered via stereotaxic injection into the ventricles for whole-brain distribution. A separate cohort of 5xFAD mice received daily oral gavage of glucosamine of 457 mg/kg/day, with the dose calculated based on human therapeutic equivalent dose of ∼2500mg/day or 36 mg/kg/day^52^ followed by standard human to mouse dose calculation^53^. Following a two-week intervention period, we assessed glycan levels using spatial glycomics and performed neurobehavioral testing to evaluate cognitive outcomes. Heatmap analysis revealed that shPGM3 treatment resulted in a global reduction in brain glycosylation levels (Fig. 5D), with spatial glycomics confirming decreases in representative N-glycans across cortical and hippocampal regions (Fig. 5E). Conversely, glucosamine supplementation significantly increased glycosylation across the brain, as shown by global heatmap analysis (Fig. 5F) and representative spatial mapping of key glycans (Fig. 5G).

To assess the functional impact of these treatments, we performed a social memory test, a behavioral paradigm that measures recognition memory by tracking social interaction time across repeated exposures^54^. In this test, mice are expected to progressively reduce their interaction time with a familiar conspecific over four trials, reflecting intact social recognition memory (Fig. 5H). Notably, shPGM3-treated 5xFAD mice exhibited significant improvement in social memory, with interaction patterns when compared to the shScr treated mice (Fig. 5I). It is worth noting that the injection alone would impact basal memory of the 5xFAD mice. In contrast, glucosamine-treated 5xFAD mice exhibited a further exacerbation of social memory deficits, showing no recognition of previously encountered conspecifics across trials (Fig. 5J). To further investigate neuropathological changes, we performed immunofluorescence analysis of reactive astrocytes and Aβ staining to quantify β-amyloid plaques (Extended Fig. 8). Interestingly, neither shPGM3 nor glucosamine treatment impacted the number of reactive astrocytes or the abundance of β-amyloid plaques during the duration of the experiment while resulting in behavioral benefit (Extended Fig. 8). These findings demonstrate that glycosylation is an actionable target for AD-related neurocognitive deficits. The improvement in social memory following glycan biosynthesis inhibition suggests that hyper-glycosylation contributes functionally to AD-related behavioral impairments. Conversely, the worsening of social memory with glucosamine supplementation supports the notion that elevated glycan biosynthesis exacerbates disease pathology. Collectively, these results establish a causal role for hyper-glycosylation in AD-associated cognitive dysfunction and highlight the therapeutic potential of targeting glycan metabolism to mitigate neurodegenerative progression.

### Real-World Impact of Glucosamine in the Dementia Patient Population

Given that glucosamine exacerbates cognitive deficits in the 5xFAD mouse model, we sought to determine whether this effect extends to human populations. As glucosamine is a widely available over-the-counter supplement commonly used for joint health, it is probable that some dementia patients utilize it regularly. To investigate whether glucosamine usage influences clinical outcomes in AD, we analyzed clinical records from the University of Florida Health (UF Health) system, identifying over 50,000 patients diagnosed with AD or Alzheimer’s disease-related dementias (ADRD). Using physician notes and prescription records, we identified patients who had documented glucosamine usage for at least one year following a dementia diagnosis. Mild cognitive impairment (MCI), clinically defined as a prodromal phase rather than a definitive dementia diagnosis, served as a practical pre-dementia comparison group. Further, since cognitively healthy individuals tend to have fewer healthcare visits, MCI group provides a more appropriate comparator group for those with confirmed dementia (Fig. 6A). In addition to AD and MCI, we also considered patients with ADRD—which includes AD, vascular dementia (VD), Lewy body dementia (LBD), and frontotemporal dementia (FD)—as a separate group. After applying inclusion criteria, we identified 9,408 AD patients, 15,073 ADRD patients, and 41,884 MCI patients for survival analysis (supplemental Fig. 6-8).

To ensure the accuracy and consistency of the glucosamine exposure classification, we first assessed the percentage of patients within each diagnostic category (AD, ADRD, and MCI) who had documented glucosamine usage. Across all three groups, approximately 8% of patients were identified as glucosamine users, confirming that physician notes provide a reproducible and reliable method for classifying supplement use (Fig. 6B). Next, we performed a 10-year survival analysis in all three cohorts, adjusting for age (≥ 50 years old). Since MCI patients are generally younger at diagnosis compared to AD and ADRD patients, age adjustment was necessary to account for baseline survival differences (supplemental Fig. 8). Our analysis revealed that glucosamine usage was associated with a 25% increase in mortality risk among AD patients (p = 0.0023, Fig. 6C) and a 35% increase in mortality risk among ADRD patients (p < 0.0001, Fig. 6D). While both sexes are negatively impacted, there are some slight variations observed between male and female patients (supplemental Fig. 6-7). In contrast, glucosamine usage did not significantly impact mortality risk in the MCI cohort after age adjustment (Fig. 6E), suggesting that the effects of glucosamine may be specific to individuals with established neurodegeneration rather than the general aging population.

### Glucosamine Supplementation in Wild-Type Mice

Given that glucosamine supplementation did not impact survival in the MCI patient cohort, we sought to determine whether the absence of an effect in this population reflects an inherent resilience of the non-diseased brain to glucosamine-induced hyper-glycosylation. To address this question, we subjected 8-month-old WT mice to the same glucosamine treatment regimen as used in 5xFAD mice. Mice received daily oral gavage of glucosamine for two weeks, with the dose adjusted based on human-equivalent scaling to match prior preclinical interventions (Extended Fig. 7A). Following the treatment period, we performed spatial glycomics analysis to assess glycan abundance across brain regions, followed by behavioral testing to determine whether glucosamine altered cognitive function in non-diseased mice.

Strikingly, glucosamine supplementation did not result in hyper-glycosylation in WT mouse brains (Extended Fig. 7B), in contrast to the robust increase observed in 5xFAD mice. Detailed glycomics analysis revealed no significant changes in N-glycan abundance across multiple glycan species (Extended Fig. 7C), suggesting that normal glycan homeostasis is maintained in the absence of AD pathology.

Furthermore, we assessed social memory performance using a four-trial system in which mice were exposed to the same social stimulus multiple times. In cognitively intact mice, social interaction time is expected to decrease over successive trials, reflecting recognition of the familiar conspecific (Extended Fig. 7D). Both vehicle- and glucosamine-treated WT mice exhibited this expected pattern of behavior, with progressive reductions in interaction time across trials. Importantly, there was no statistically significant difference between the two groups, indicating that glucosamine supplementation did not impair social recognition memory in WT mice (Extended Fig. 7E). This contrasts with the deficits observed in 5xFAD mice, supporting the notion that the normal brain possesses intrinsic resilience mechanisms that buffer against the metabolic perturbations induced by glucosamine supplementation.

## Discussion

MALDI-MSI, has proven to be a powerful tool for hypothesis-generating research, allowing for the spatial characterization of metabolic and molecular alterations directly in tissue samples^33,55,56^. This study utilized MALDI-MSI based spatial multiomics to uncover an hyper-glycosylation phenotype in AD brains. While such findings offer critical insights, it is essential to validate them using controlled experimental models to establish causality and mechanistic underpinnings. By employing the 5xFAD mouse model, we demonstrated that hyper-glycosylation is a conserved pathological feature of neurodegeneration and further identified increased glycan biosynthesis as the primary driver of this metabolic dysregulation. The integration of MALDI-MSI with stable isotope tracing and gene expression analysis allowed us to precisely delineate the metabolic origins of this phenotype, reinforcing the importance of spatial metabolomics in neurodegenerative disease research.

Protein glycosylation is an essential post-translational modification with profound implications for neuronal function^57^, and our findings highlight an intriguing hyper-glycosylation phenotype that emerges as a hallmark of AD. Hyper-glycosylation, driven primarily by increased biosynthesis rather than lysosomal salvage pathways, is particularly notable given the reported lysosomal pathway dysfunction in AD^58,59^. The presence of hyper-glycosylation does not invalidate evidence of lysosomal defects but instead underscores a more complex scenario involving multiple points of metabolic disruption that collectively contribute to AD pathology, warranting further investigation. Indeed, the observed upregulation of glycosylation under glucose hypo-uptake conditions suggests a compensatory metabolic mechanism aimed at mitigating broader metabolic deficits. Speculatively, this may represent an adaptive cellular response attempting to preserve neuronal function or survival by modulating protein processing, trafficking, and possibility immune interactions through enhanced glycosylation. Glycoproteomic analyses further support this hypothesis, indicating that the majority of glycosylated proteins identified in AD are neuronal membrane proteins critical for action potential propagation and synaptic transmission. Future research should thus explore these compensatory mechanisms more deeply, potentially revealing novel therapeutic targets or diagnostic markers associated with metabolic dysregulation in AD.

Targeting glycosylation as a therapeutic strategy presents a unique challenge due to the necessity of preserving central glucose metabolism, which is vital for neuronal function^47,60^. Unlike broad metabolic interventions that disrupt glucose utilization^61^, targeting the hexosamine biosynthetic pathway (HBP) offers a more selective approach^62^. Notably, our data indicate that neither glucosamine supplementation nor shRNA-mediated inhibition of PGM3 significantly impacted neuroinflammatory markers such as astrocytosis or GFAP levels, nor did they substantially alter amyloid-beta formation in the mouse model. This suggests that targeting metabolic pathways might represent a viable therapeutic strategy independent of addressing neuroinflammation^22^ or amyloid clearance^63^, or potentially as complementary to existing therapies. Supporting this perspective, a recent study demonstrated that pharmacological inhibition of IDO1 and subsequent restoration of glucose metabolism could effectively reverse cognitive symptoms in multiple mouse models of AD without directly affecting hallmark AD pathology^9^. Our findings reinforce this concept by highlighting PGM3 as a selective target, with its inhibition reducing glycan biosynthesis while improving cognitive memory. While no blood-brain barrier-permeable inhibitors of PGM3 currently exist, the identification of such compounds represents an important next step. Future studies should focus on developing and testing small-molecule inhibitors that selectively reduce pathological glycosylation in preclinical AD models, paving the way for potential clinical translation.

Glucosamine is a widely used dietary supplement that is not subject to FDA regulation, and can vary considerably in quality and purity^64^. Our data suggest that it does not negatively impact cognitively normal individuals but significantly worsens outcomes in AD patients. Across our real-world EHR analysis^65^, glucosamine use was associated with a nearly 35% increase in mortality in both AD and ADRD populations. It is important to consider this population level data with human brain specimens and mouse AD models through genetic and dietary challenges. Together, they make a compelling argument that glucosamine as a negative driver of neurodegeneration. Interestingly, glucosamine usage did not lead to an increased mortality rate in the MCI population as a whole, a group that is clinically not classified as having dementia. However, it is important to acknowledge that in real-world clinical settings, MCI is often used as a syndromic diagnosis. In the past, many neurologists coded AD or dementia based on clinical presentation rather than underlying pathology. As a result, patients diagnosed with MCI may, in fact, have underlying AD. This diagnostic ambiguity underscores the need to interpret MCI-specific findings with caution, as they may reflect a heterogeneous population with varying degrees of neurodegenerative progression.

This suggests that the adverse effects of glucosamine may be contingent on an underlying neurometabolic state. Due to this phenotype-specific vulnerability, we proceeded to evaluate the impact of glucosamine supplementation in wild-type mice as a model of the metabolically normal brain. As observed in the MCI patient population, glucosamine-treated WT mice did not exhibit elevated N-glycan levels nor impairments in social memory performance. These data collectively highlight a crucial distinction between the metabolic homeostasis of the normal brain and the dysregulated glycan biosynthetic machinery present in dementia. Our findings suggest that hyper-glycosylation and the deleterious effects of glucosamine are not generalizable across all individuals but are uniquely positioned to exacerbate metabolic vulnerability and cognitive decline in the context of neurodegeneration. In the United States, an estimated 6.7 million individuals are currently living with AD and another 7 million individuals with ADRD. Given our calculation that approximately 8% of patients in our dataset were using glucosamine, this suggests that over 1 million patients may be unknowingly exacerbating their disease progression through glucosamine supplementation. These findings emphasize the urgent need for a well-designed, double-blind clinical trial to systematically evaluate the impact of glucosamine on cognitive decline in dementia patients. Establishing a clear clinical framework for evaluating its effects in at-risk populations is crucial to determining future best practices.

One potential concern is the high prevalence of osteoarthritis (OA) and rheumatoid arthritis (RA) among glucosamine users. Recent randomized doubled blinded, and placebo-controlled trails have demonstrated that glucosamine has limited or no efficacy in alleviating joint pain associated with OA or RA^66,67^. In fact, most OA and RA patients in clinical practice rely on non-steroidal anti-inflammatory drugs (NSAIDs) or disease-modifying therapies for symptom management^68,69^, rather than glucosamine. Additionally, glucosamine is not recommended by the U.S bone and joint initiative as a treatment for joint pain^70^.

Furthermore, current marketing strategies for glucosamine products overwhelmingly target physically active and healthy individuals seeking to preserve joint mobility and flexibility, rather than patients experiencing chronic arthritic pain. Therefore, we do not believe OA or RA represent meaningful covariates in our population-level study. Additionally, the two UK Biobank study reported different trends of AD diagnosis rates in glucosamine users, one study suggested a lower rate of AD with long term glucosamine use^71^, and another reported no difference from the same cohort of patients in AD and ADRD, but lower risk for vascular dementia^72^. Importantly, our findings do not conflict with these data, as our study specifically examines patients who already have AD rather than the general population at risk of developing the disease, i.e. incidence of diagnosis. The possibility remains that glucosamine may be beneficial for cognitively normal individuals while exacerbating pathology in those with established neurodegeneration. Based on our discussion above, this difference critically highlights the need for rigorous controlled clinical studies to delineate the differential longitudinal effect of glucosamine supplementation in different groups.

This study underscores the broader potential of MALDI imaging and spatial biology in AD research. Incorporating cell-type-specific spatial information through emerging technologies such as CODEX, MIBI-TOF, or single-cell spatial transcriptomics will further refine our understanding of how metabolic dysregulation intersects with cellular pathology. We expect hyper-glycosylation to separately impact neuronal function^73^, glial response^74^, blood brain barrier^23^, glymphatic systems^75^. Additionally, the development of pharmacological interventions targeting the hexosamine pathway in animal models represents a promising avenue for therapeutic exploration. Finally, given our findings on glucosamine supplementation, we strongly advocate for a large-scale, double-blind clinical trial to definitively determine its impact on AD progression at the population level. This study provides a foundation for future translational efforts aimed at modulating glycan metabolism as a therapeutic approach in AD.

## Acknowledgments

This study was supported by National Institute of Health (NIH) grants R01AG066653, R01CA266004, R01AG078702, R01CA288696, RM1NS133593 to R.C.S., R35NS116824 to M.S.G., R35GM142701 to L.C, T32HL134621 to H.A.C. D.B.A and supported in part by an NIH award, S10 OD030293. Z.L is supported by the MBI Gator NeuroScholar Program. The UF Neuromedicine Human Brain and Tissue bank is supported by the McKnight Brain Institute, the Center for Translational Research in Neurodegenerative diseases at the University of Florida and NIH grant P30AG066506. S.P. is the Charlotte and Howard Zimmerman Rising Star Professor with the Norman Fixel Institute for Neurological Diseases. Large Language Models, e.g. ChatGPT, was used to make minor grammatical improvements in the text.

## Author contribution statement

Conceptualization, R.C.S.; Methodology, R.C.S.; Investigation, T.R.H., Z.L., T.M.M., R.S.N., R.A.R., H.A.C., X.M., A.C.M., A.F.P., C.M.S., J.S., F.C., A.R.C., M.G.C., C.J.S., L.E.A.Y., D.D.K., C.L., A.M.R., P.A., R.J.D., S.P., Y.H., X.H., J.B., S.P., C.W.V.K., M.S.G., R.C.S.; Writing – Original Draft, R.C.S.; Writing – Review & Editing, R.C.S., R.J.D., C.W.V.K., M.S.G.; Funding Acquisition, M.S.G., R.C.S.; Resources, M.S.G., R.C.S.; Supervision, M.S.G., R.C.S.

## Methods

### Chemicals, reagents, antibodies, and cell lines

High-performance liquid chromatography (HPLC)-grade acetonitrile, ethanol, methanol, water, trifluoroacetic acid (TFA), N-(1-Naphthyl) ethylenediamine dihydrochloride (NEDC), and recombinant isoamylase were purchased from Sigma-Aldrich. α-cyano-4-hydroxycinnamic acid (CHCA) matrix was purchased from Cayman Chemical. Histological-grade xylenes were purchased from Spectrum Chemical. Citraconic anhydride for antigen retrieval was obtained from Thermo Fisher Scientific. Recombinant PNGaseF Prime was obtained from N-Zyme Scientifics (Doylestown, PA, USA). Bruker IntelliSlides (Bruker Daltonics).

### Human and Mouse Tissue Sources and Preparation

De-identified human brain tissue samples were obtained under IRB-approved protocols from the University of Florida Alzheimer’s Disease Research Center (UF-ADRC). Both fresh frozen and formalin-fixed, paraffin-embedded (FFPE) frontal cortex specimens from cognitively normal and Alzheimer’s disease individuals were utilized. Post-mortem intervals (PMIs), age, and sex were matched between groups, and Braak staging was conducted for all AD cases. Frozen tissue blocks were cryosectioned at 10 µm thickness and mounted onto indium tin oxide (ITO)-coated conductive slides for downstream MALDI imaging and molecular analyses. Mice were housed in climate-controlled environment with a 14 (light)/10 (dark) hours light/dark cycle with temperature (18-23°C) and humidity (50-60%) control. Water and solid diet provided *ad libitum* throughout the study (Tekad #2018). The University of Florida Institutional Animal Care and Use Committee has approved the animal procedures under the protocol number IACUC202200000586. Mouse brain tissue was obtained from 5xFAD transgenic mice and wild-type (WT) littermate controls maintained under standard housing conditions with ad libitum access to food and water (JAX stock #008730). Female mice aged 8–9 months were used for all experiments to reflect a time point of established AD-like pathology. For stable isotope tracing experiments, mice received a 13C-labeled liquid diet for 24 hours (pulse group) or a 24-hour pulse followed by a 48-hour washout period (chase group). In separate cohorts, mice were treated with glucosamine sulfate by oral gavage at 457 mg/kg/day for two weeks. All animal protocols were approved by the University of Florida Institutional Animal Care and Use Committee (IACUC).

### Lentiviral Delivery and Stereotaxic Injections

To knock down glycan biosynthesis, short hairpin RNA targeting PGM3 (shPGM3, TL505905V; Origene) or control shScrambled lentivirus (TR30021V; Origene) was bilaterally injected into the lateral ventricles of 5xFAD mice. Injections were performed using a stereotaxic frame at coordinates relative to bregma (AP: -0.2 mm; ML: ±1.0 mm; DV: -2.2 mm). Lentiviral particles (1.0 × 10^9 TU/ml) were delivered in a 2 µL volume (at a rate of 0.5 µL/min) per ventricle using a Hamilton syringe and allowed to diffuse for 5 minutes post-injection before needle withdrawal.

### Sample Collection and Preparation

Mice were euthanized by cervical dislocation and decapitation. Immediately following euthanasia, the brains were surgically resected within 30 seconds. Mouse brains were dissected into two hemispheres coronally to expose regions to be scanned by MALDI. The brain tissue was first rinsed in 1x Phosphate-Buffered Saline (PBS) and subsequently rinsed twice with deionized water. The rinsed tissues were blotted dry, then slow-frozen over isopentane chilled with dry-ice for seven minutes, according to previously established protocols^69,70^, to ensure tissue stability and optimal preservation of analytes. Post-freezing, the samples were stored at -80°C until further processing.

### MALDI Imaging for Spatial Metabolomics, Lipidomics, and Glycomics

Frozen brain tissues were sectioned at −15°C using a Leica CM1860 cryostat and mounted onto a frozen chuck with OCT compound. Coronal sections of 10 µm thickness were collected, thaw-mounted onto Bruker IntelliSlides, and stored at −80°C. Prior to matrix application, slides were fixed by vacuum desiccation for one hour. Each slide was sequentially processed for spatial metabolomics, lipidomics, glycomics, and glycogen imaging. For metabolomics and lipidomics, N-(1-Naphthyl) ethylenediamine dihydrochloride (NEDC) matrix was applied using an HTX M5 sprayer with 14 passes of 7 mg/mL NEDC in 70% methanol at 30°C and 10 psi, with a spray rate of 0.06 mL/min and tray heated to 50°C. Following metabolite and lipid imaging, slides were washed with ice-cold 100% methanol for 5 minutes to remove NEDC and fixed overnight in 10% neutral buffered formalin. For glycomics and glycogen analysis, mouse tissue sections were processed through ethanol and water rinses, followed by antigen retrieval in citraconic anhydride buffer (25 µL in 50 mL H2O, pH 3) at 95°C for 25 minutes. For human brain FFPE sections, lipid removal was performed using three sequential 1-hour washes in xylene prior to enzymatic treatment. Enzyme application was performed using HTX sprayer with a 0.2 mL solution containing isoamylase (3 U/slide) and PNGase F (20 µg/slide) at 45°C and 900 mm/min spray velocity. Slides were incubated at 37°C for 2 hours in a humidified chamber and then vacuum desiccated. CHCA matrix (0.04 g in 5.7 mL 50% acetonitrile/50% water with 5.7 µL 25% TFA) was sprayed prior to imaging. MALDI mass spectrometry imaging was performed on a Bruker timsTOF fleX using a 46 µm × 46 µm laser raster to produce 50 µm × 50 µm pixels. Image acquisition and setup were performed using the autopilot feature in flexImaging v6.0 to minimize user-dependent variability. A standardized flexImaging.mis file was generated from scanned high-resolution .tif images and applied to all omics runs to ensure consistent region masking and pixel alignment. For spatial metabolomics, imaging was conducted in negative ion mode with the following settings: MS1 scan, m/z 20–750, 90% laser power, 1 burst of 396 shots, 10,000 Hz frequency, 30 V MALDI plate offset, −60 V deflection 1 delta, 200 Vpp Funnel 1 RF, 200 Vpp Funnel 2 RF, 200 Vpp Multipole RF, 7 eV collision energy, and 700 Vpp collision RF. Lipidomics imaging was also performed in negative ion mode immediately following metabolomics, with MS1 scan settings of m/z 300–2000, 80% laser power, 1 burst of 300 shots, and the same raster, frequency, and transfer parameters as metabolomics. Glycomic imaging followed glycan preparation and was performed in positive ion mode with MS1 scan settings of m/z 700–4000, 37% laser power, 1 burst of 320 shots, 10,000 Hz frequency, 50 V MALDI plate offset, 70 V deflection 1 delta, 500 Vpp Funnel 1 RF, 500 Vpp Funnel 2 RF, 500 Vpp Multipole RF, 10 eV collision energy, and 4000 Vpp collision RF. Post-acquisition data were aligned and analyzed using SCiLS Lab software to extract spatially resolved ion distributions and intensities.

### Stable Isotope Tracing and Ion Mobility Mass Spectrometry

To assess glycan biosynthetic flux, we employed a stable isotope tracing protocol using a 13C6-glucose-enriched liquid diet. Tissue sections from pulse and chase cohorts were analyzed using matrix-assisted laser desorption/ionization (MALDI) mass spectrometry coupled with ion mobility separation to resolve isotopically labeled glycans. For trapped ion mobility spectrometry (TIMS), data were acquired on a Bruker timsTOF fleX as described above, with isotopologue separation achieved through collisional cross section (CCS) measurements. To further resolve and image glycogen and N-glycan isotopologues, selected tissue sections were analyzed using a Waters Synapt G2-Si high-definition mass spectrometer equipped with traveling wave ion mobility (TWIM). The instrument operated in positive ion mode over an m/z range of 500–3,000. A laser frequency of 1,000 Hz, energy of 200 arbitrary units (AU), and laser spot size of 75 µm were used. Ion mobility settings followed established protocols, with a trap entrance energy of 2 V, trap bias of 85 V, and DC/exit of 0 V. Wave velocity settings were configured as follows: trap 9.6 m/s, IMS 4.6 m/s, and transfer 17.4 m/s. Wave height settings included trap 4 V, IMS 42.7 V, and transfer 4 V, with a variable wave ramp down of 1,400 m/s. Data acquisition was performed using MassLynx v4.2, and images were generated with HDI software v1.5 (Waters Corp), utilizing the built-in peak integration function to correct for mass drift over the course of acquisition. All images were normalized to total ion current (TIC) on a per-pixel basis.

### LC-MS Analysis of UDP-Sugar Precursors

Tissue samples were homogenized at 4°C in 80% LC-MS grade acetonitrile aqueous solution. Homogenates were centrifuged two to three times at maximum speed to remove cellular debris, and supernatants were collected into LC-MS vials. Samples were randomized and analyzed in a blinded manner using a 1290 UHPLC system coupled to a 6550 iFunnel Q-TOF mass spectrometer (Agilent Technologies), operated with MassHunter software v10.1. Chromatographic separation was performed on a SeQuant ZIC-pHILIC column (150 × 2.1 mm, 5 µm, PEEK-coated). The mobile phases were 10 mM ammonium acetate in water, adjusted to pH 9.8 with concentrated NH4OH (solvent A), and acetonitrile (solvent B). The elution gradient was as follows: 0–15 min linear gradient from 90% to 30% B; 15–18 min at 30% B; 18–19 min linear gradient from 30% to 90% B; followed by 9 min re-equilibration at 90% B. The flow rate was 0.25 mL/min with an injection volume of 20 µL, and the column temperature was maintained at 25°C. ESI source parameters were: dry gas 225°C at 18 L/min, sheath gas 350°C at 12 L/min, fragmentor voltage 175 V, nozzle voltage 500 V, and capillary voltage ±3,500 V for positive and negative modes. Full scan data were acquired across an m/z range of 40–1,700 at 1 spectrum/s in profile mode. Data were processed using Profinder B.08.00 SP3 (Agilent Technologies) against an in-house library of authentic standards, with matching criteria set at 10 ppm mass tolerance and 0.5 min retention time tolerance. All peak integrations were manually reviewed for consistency.

### RT-PCR and Digital PCR for Glycosylation Enzymes

Total RNA was extracted from frozen brain tissue using a conventional protocol involving TRIzol reagent and chloroform separation, followed by overnight precipitation in isopropanol at −20°C. The following day, DNA was denatured using phenol, and RNA pellets were washed, cleaned, and resuspended in 10 mM Tris-HCl buffer (pH 7.5). RNA quality was assessed using a Nanodrop spectrophotometer, and aliquots (0.5–1.5 µg) were stored in single-use tubes at −80°C for no longer than one week before downstream analysis. Complementary DNA (cDNA) was synthesized using the iScript One-Step RT-PCR kit with reverse transcriptase (Bio-Rad) according to the manufacturer’s instructions. Thermal cycling conditions included priming at 25°C for 5 minutes, reverse transcription at 46°C for 20 minutes, and enzyme inactivation at 95°C for 1 minute, followed by a 4°C hold. Gene expression was quantified using commercially available 96-well RT-PCR arrays targeting glycosylation-related genes (Bio-Rad Glycosylation SAB Target Lists H96 and M96 for human and mouse, respectively).

### Glycoproteomics Analysis

Brain tissue samples were subjected to membrane protein extraction by homogenization in high salt buffer (2.0 M NaCl, 5.0 mM EDTA, pH 7.4, supplemented with protease inhibitors), followed by repeated passage through a 20-gauge needle and probe sonication. Lysates were centrifuged at 45,000 rpm for 15 minutes at 4°C. Pellets were resuspended in sodium carbonate buffer (0.1 M Na2CO3, 1.0 mM EDTA, pH 11.3) and incubated on ice for 30 minutes before a second ultracentrifugation at 45,000 rpm for 45 minutes. Membrane-enriched pellets were dissolved in urea lysis buffer (8.0 M urea, 1.0 M NaCl, 4% CHAPS, 100 mM DTT, 200 mM Tris HCl, pH 8.0) and denatured at 50°C for 45 minutes. Samples were then alkylated with iodoacetamide (IAA, 55 mg) in the dark for 45 minutes at room temperature. Proteins were precipitated using chloroform/methanol/water extraction and resuspended in 50 mM ammonium bicarbonate (pH 8.0). Protein concentrations were quantified by BCA assay, and 100 µg of membrane protein was digested with trypsin (1:20) at 37°C for 24 hours. Glycopeptides were analyzed on an Ultimate 3000 RSLCnano system coupled to a Thermo Eclipse mass spectrometer. Separation was achieved using 15 cm × 75 µm C18 nanoLC columns with 3 µm particles, employing a 180-minute gradient from low to high acetonitrile in 0.1% formic acid. MS1 spectra were acquired at 120k resolution in the Orbitrap, and precursor ions were selected for HCD fragmentation over 3-second cycles. Precursor ions with charge states of +1 or unknown were excluded, and dynamic exclusion was set to 30 seconds.

Fragment ions were acquired at 30k resolution. EThcD was triggered based on detection of the HexNAc oxonium ion (m/z 204). Data were analyzed using Proteome Discoverer v3.0. Peptides were identified using CHIMERYS search with a sum PEP score >15 for global proteomics. Glycopeptides were identified using the Byonic node with precursor and fragment mass tolerances of 5 ppm and 10 ppm, respectively. Variable modifications included deamidation (N/Q), oxidation (M), and carboxymethylation (C). Glycopeptide identifications were filtered based on log probability >1 and ppm error <3. N-glycan libraries included “N-glycan 182 human no multiple fucose” for human samples and “N-glycan 309 mammalian no sodium” for mouse samples. O-glycan identification was performed using the “9-Most-common glycans” library for both species. CHIMERYS and Byonic output files were curated and shared with study collaborators for independent verification.

### Behavioral Testing: Social Memory Assay

Social recognition memory was assessed in the animal facility at the University of Florida using a previously established protocol. Mice were acclimated to the behavior room for 30 minutes and subsequently placed in a clean rat cage, divided by a cardboard partition, for an additional 10-minute habituation period. Experimental mice were randomly assigned to testing order and introduced to the same novel conspecific once daily over five consecutive days, with trials separated by 24-hour intervals. Behavioral interactions were recorded during each session and quantified using DeepLabCut, an open-source deep learning platform for pose estimation. Frame-by-frame analysis was used to quantify the duration the experimental mouse spent within 1 cm of the novel conspecific during each trial.

### Immunofluorescence and Immunohistochemistry Analysis

Brain tissue sections from mouse and human samples were fixed in 10% neutral-buffered formalin (NBF), paraffin-embedded, and stored prior to sectioning. Mice were sacrificed via cervical dislocation, and tissues were resected and fixed immediately. Human tissue samples were obtained from the University of Kentucky Biospecimen Facility under approved protocols. Paraffin-embedded sections (4 μm) were used for immunohistochemistry. Slides were dewaxed, rehydrated, and subjected to antigen retrieval using Ventana CC2 buffer at 37°C for 1 hour. Primary antibodies included GFAP (astrocyte marker; 1:100, GTX85454; GeneTex), 6E10 (Aβ plaque marker; 1:100, 803015; Biolegend), Laforin (1:100, ab129321; Abcam), GYS1 (1:100, #3886, Cell Signaling), and PYGB (1:100, 12075-1-AP; Proteintech). Primary and secondary antibody incubations were carried out at room temperature for 60 minutes each. H&E staining was conducted using standard protocols with hematoxylin, eosin, graded ethanol dehydration, and coverslipping. Digital slide images were acquired using a ZEISS Axio Scan.Z1 platform. Quantitative image analysis was performed using HALO software (Indica Labs) with multiplex IHC and tissue microarray modules to assess marker expression and pathology across matched regions of interest.

### Human EHR-Based Cohort Identification and Glucosamine Stratification

We performed a retrospective cohort study using electronic health record (EHR) data from 2012 to 2024 obtained through the University of Florida Health Integrated Data Repository (UF IDR), a centralized enterprise data warehouse integrating clinical data across UF Health systems. The UF IDR includes over billion observational facts from more than 2 million patients. The study was reviewed and granted exempt status by the University of Florida Institutional Review Board (IRB202001888). The study population included individuals with a diagnosis of mild cognitive impairment (MCI) or Alzheimer’s disease and related dementias (AD/ADRD), including AD, vascular dementia (VD), Lewy body dementia (LBD), and frontotemporal dementia (FD), identified by ICD-9-CM and ICD-10-CM codes (Supplementary Table 1). Glucosamine users were identified through a natural language processing (NLP) pipeline that extracted glucosamine-related records from clinical notes using a regex-based keyword matching approach. Notes containing relevant glucosamine terms were flagged as exposures.

### Survival and Transition Analyses from EHR Data

Data preprocessing included exclusion of records with inconsistencies, such as death dates preceding birth or diagnosis. Patients were categorized into five groups: AD, MCI, ADRD, MCI-to-AD, and MCI-to-ADRD. For groups 4 and 5, only patients with an MCI diagnosis preceding AD/ADRD were included. Within each group, patients were further classified into glucosamine users (cases) and non-users (controls). Kaplan-Meier analyses were performed to estimate survival probabilities over a 15-year window in glucosamine users versus non-users, with subgroup analyses conducted for all ages and those aged >50 years. Transitions from MCI to AD or ADRD were calculated and compared between groups. Analyses were implemented in Python (v3.10.8) using NumPy (v1.25.2), pandas (v2.2.2), and the lifelines library (v0.29).

### Quantification and Statistical Analysis

All animals and data points are included for statistical analysis. Data distribution was assumed to be normal but this was not formally tested. Statistical analyses were carried out using GraphPad Prism. All numerical data are presented as mean ± S.E.M Column analysis was performed using one-way or two-way ANOVA or Student’s t-test. P-value is indicated on the graph. The statistical parameters for each experiment can be found in the figures and figure legends.

## Extended figures

**Extended Fig. 1.**
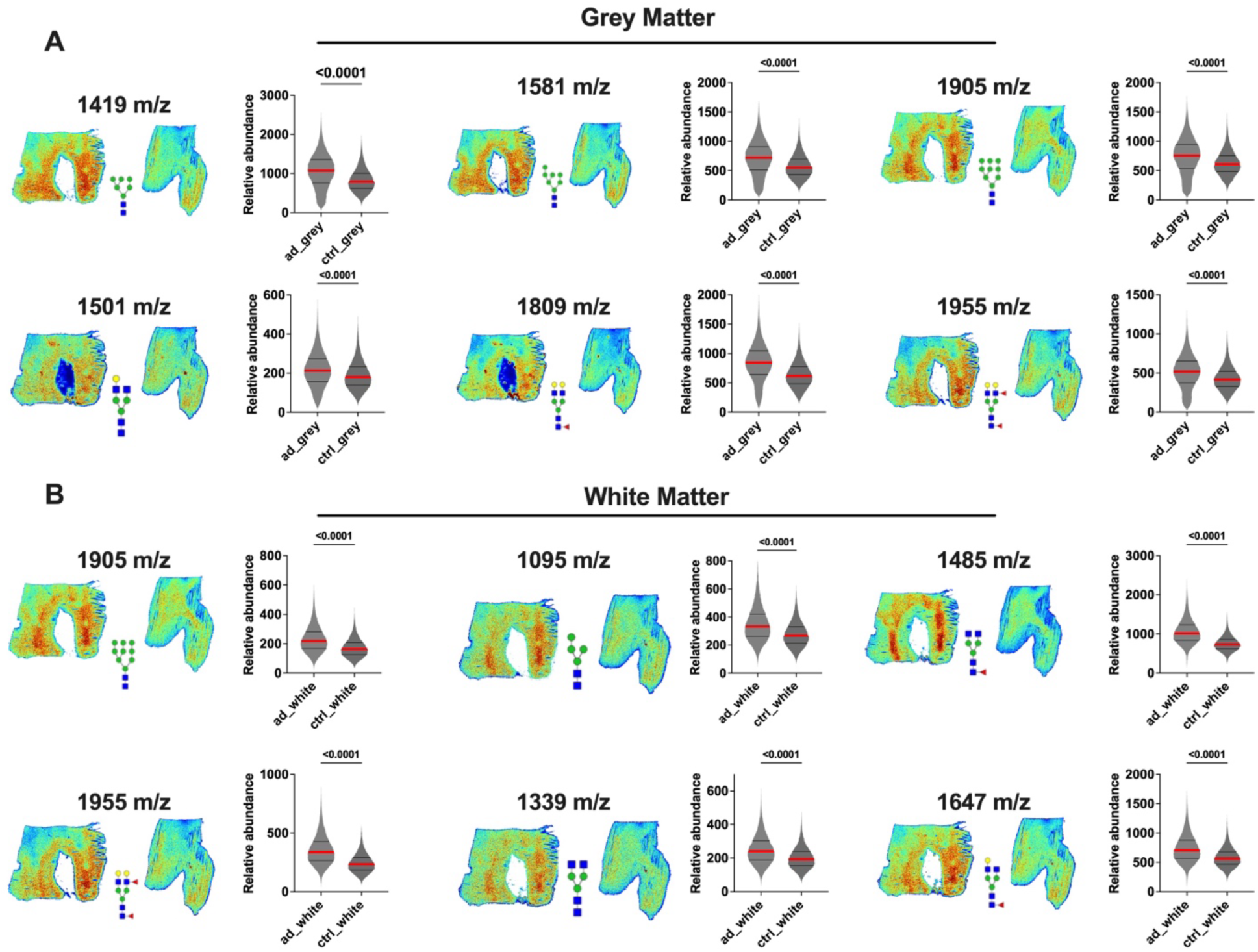
Regional glycan abundance increases in human Alzheimer’s disease brains. **a**, Spatial glycomics images and corresponding violin plots illustrating significantly elevated glycan abundance in grey matter regions of Alzheimer’s disease (AD) brains relative to controls. Glycan species (1419, 1501, 1581, 1809, 1905, and 1955 m/z) exhibit consistent and significant increases in AD. Statistical analysis performed using two-tailed t-tests; exact P-values indicated. (n>2000 pixels across three different samples) **b**, Spatial glycomics visualization and quantitative analysis of glycan abundance in white matter regions. Similar significant elevations observed across glycan species (1095, 1339, 1485, 1647, 1905, and 1955 m/z) in AD brains compared to controls (n>2000 pixels across three different samples). Statistical significance assessed by two-tailed t-tests; exact P-values indicated.

**Extended Fig. 2.**
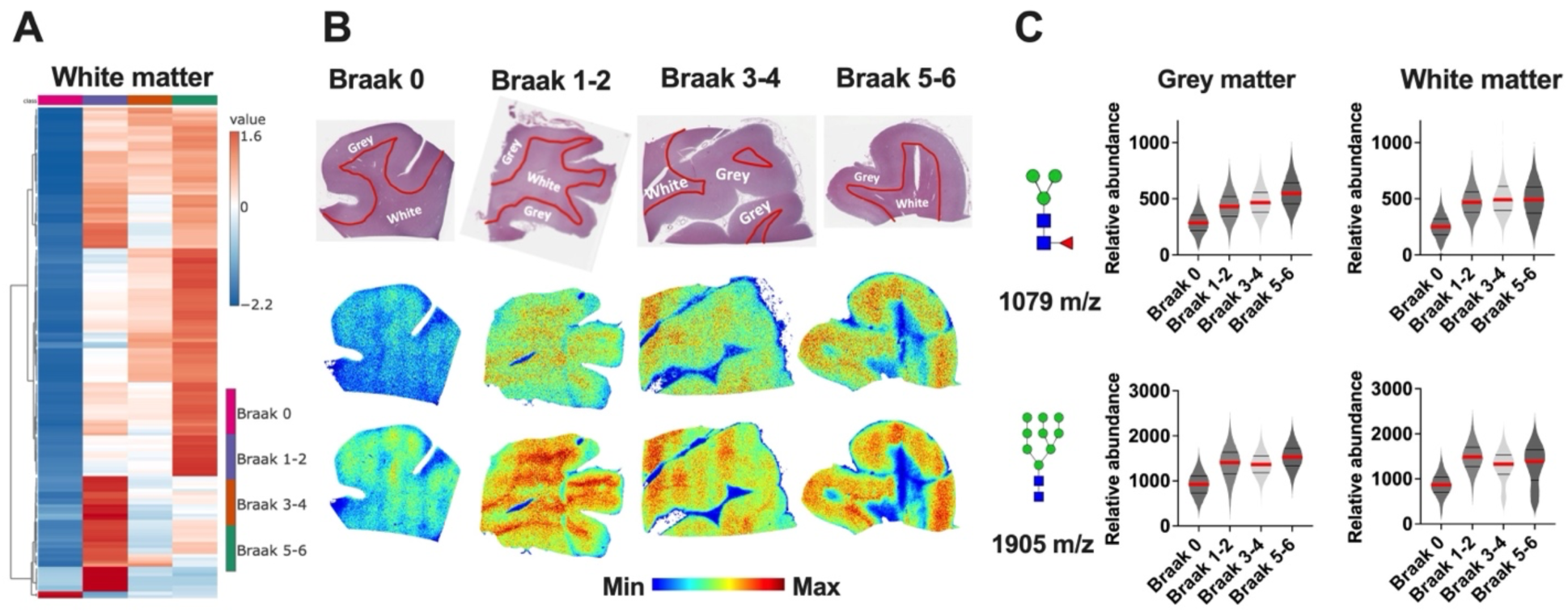
Stage-dependent glycan accumulation in white matter regions across Braak stages. **a**, Heatmap of glycan abundance in white matter across Braak stages 0, 1–2, 3–4, and 5–6, showing early-stage elevation of glycosylation in Braak 1–2 with no sustained increase in later stages. **b**, Representative histological and spatial glycomics images showing distributions of glycan species (1079 and 1905 m/z) in white matter across Braak stages. **c**, Violin plots quantifying relative abundance of 1079 and 1905 m/z glycans in both grey and white matter, revealing a transient increase in white matter glycosylation at early Braak stages without further elevation in late-stage disease.

**Extended Fig. 3.**
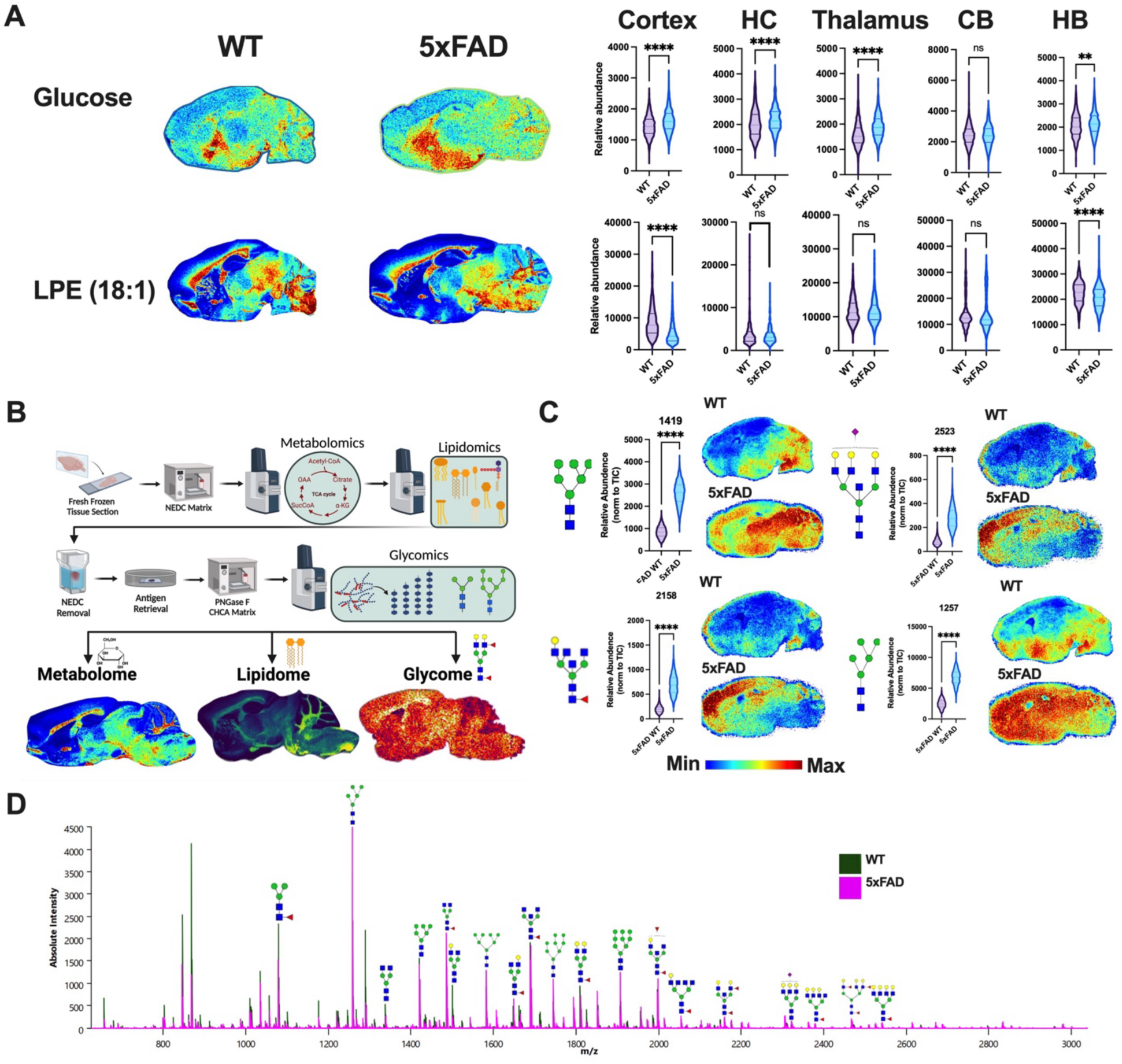
Integrated spatial multiomics confirms glycan enrichment in 5xFAD mouse brains. **a**, Spatial metabolomics and lipidomics maps of glucose and lysophosphatidylethanolamine (LPE 18:1) in wild-type (WT) and 5xFAD brains, with accompanying violin plots showing region-specific differences across cortex, hippocampus (HC), thalamus, cerebellum (CB), and hindbrain (HB). Significant reductions in glucose and elevations in LPE are observed in disease-relevant regions of 5xFAD brains (n>2000pixels across 3 biological replicates each). Statistical significance assessed by two-tailed t-tests; **b**, Schematic of spatial multiomics workflow detailing integration of NEDC-based metabolomics and lipidomics with CHCA-PNGase-based glycomics from adjacent sections. **c**, Spatial glycomics images and violin plots of representative glycan species (1419, 2158, 2523, 1237 m/z) confirm significant glycan accumulation in 5xFAD compared to WT across brain regions. Two-tailed t-tests used for comparisons. **d**, MALDI mass spectra showing elevated glycan peak intensities across a wide m/z range in 5xFAD (pink spectra) compared to WT (green spectra) mouse brains, confirming broad glycan upregulation.

**Extended Fig. 4.**
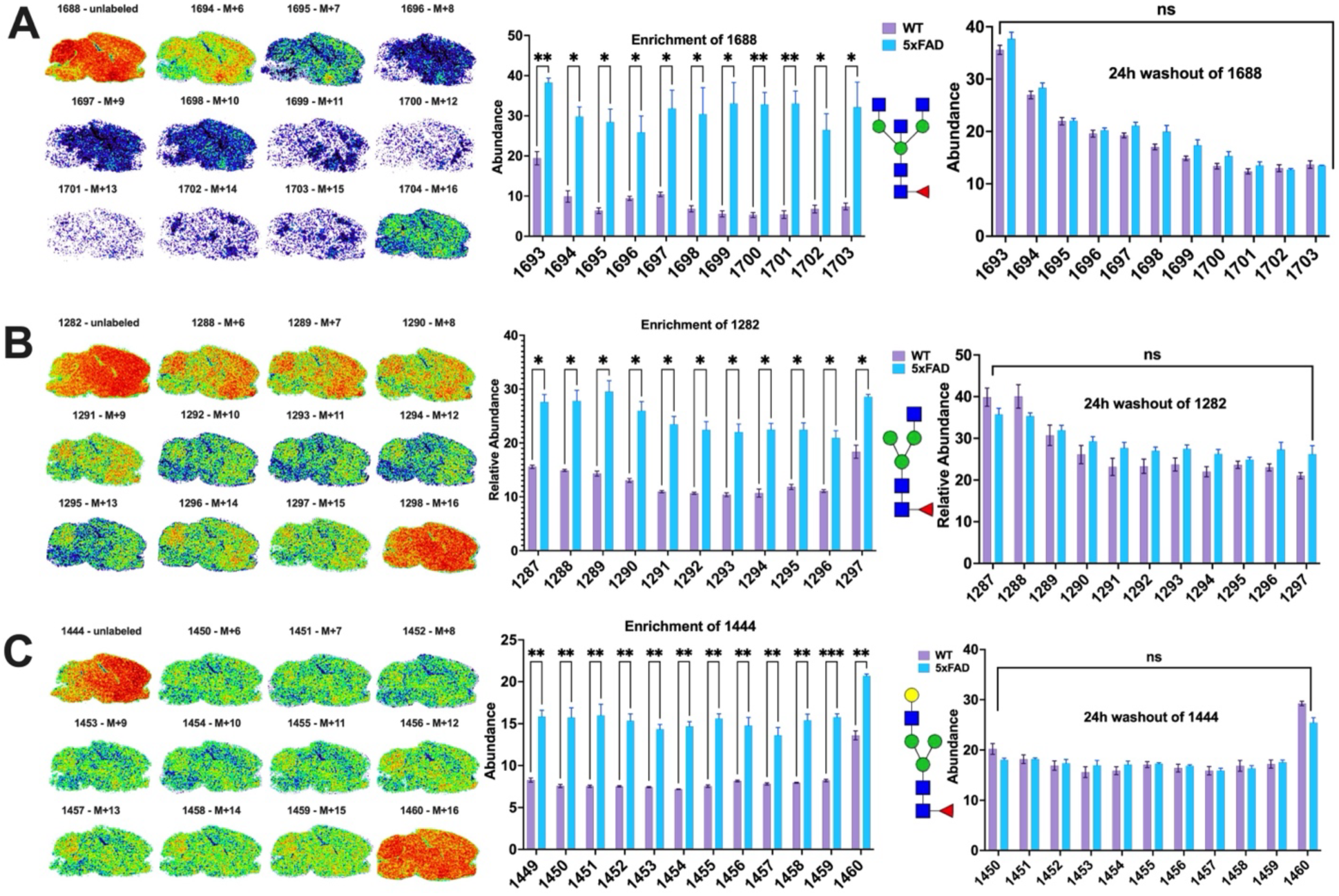
Multiple glycans exhibit increased biosynthesis but unchanged turnover in 5xFAD mouse brains. **a**, Spatial distribution and quantification of 13C6-labeled isotopologues of glycan 1688 m/z following isotope pulse in WT and 5xFAD mice. Bar graphs show significantly increased 13C incorporation across isotopologues in 5xFAD. Washout after 24 hours reveals no significant difference in turnover between genotypes. Statistical analysis via two-tailed t-tests (n=3 biological replicates) **b**, Enrichment and washout analysis of glycan 1282 m/z demonstrates elevated biosynthesis in 5xFAD with no change in degradation rates. **c**, Similar enrichment profile observed for glycan 1444 m/z, showing significantly increased labeling in 5xFAD without altered clearance.

**Extended Fig. 5.**
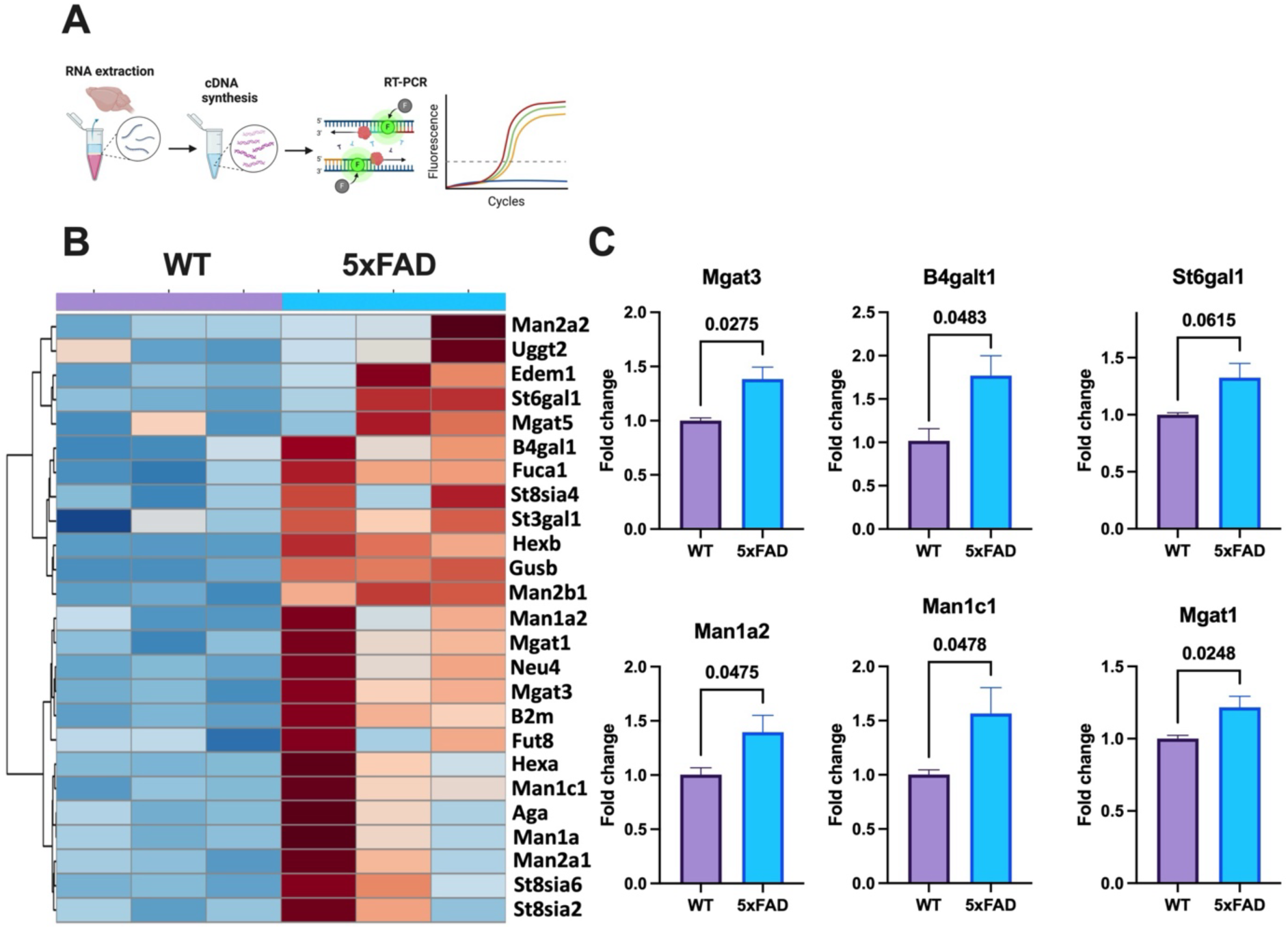
Transcriptional upregulation of glycosylation-related genes in 5xFAD mouse brains. **a**, Schematic of experimental workflow for RNA extraction, cDNA synthesis, and RT-qPCR analysis of glycosylation-associated genes. **b**, Heatmap showing gene expression profiles of 27 glycosylation-related genes in wild-type (WT) and 5xFAD mouse brains. Hierarchical clustering reveals widespread transcriptional upregulation in 5xFAD mice. **c**, Bar graphs displaying fold change comparisons for selected glycosylation-related genes (Mgat3, B4galt1, St6gal1, Man1a2, Man1c1, and Mgat1) between WT and 5xFAD. Statistical significance determined by two-tailed t-tests; exact P-values indicated; (N= 3 biological replicates).

**Extended Fig. 6.**
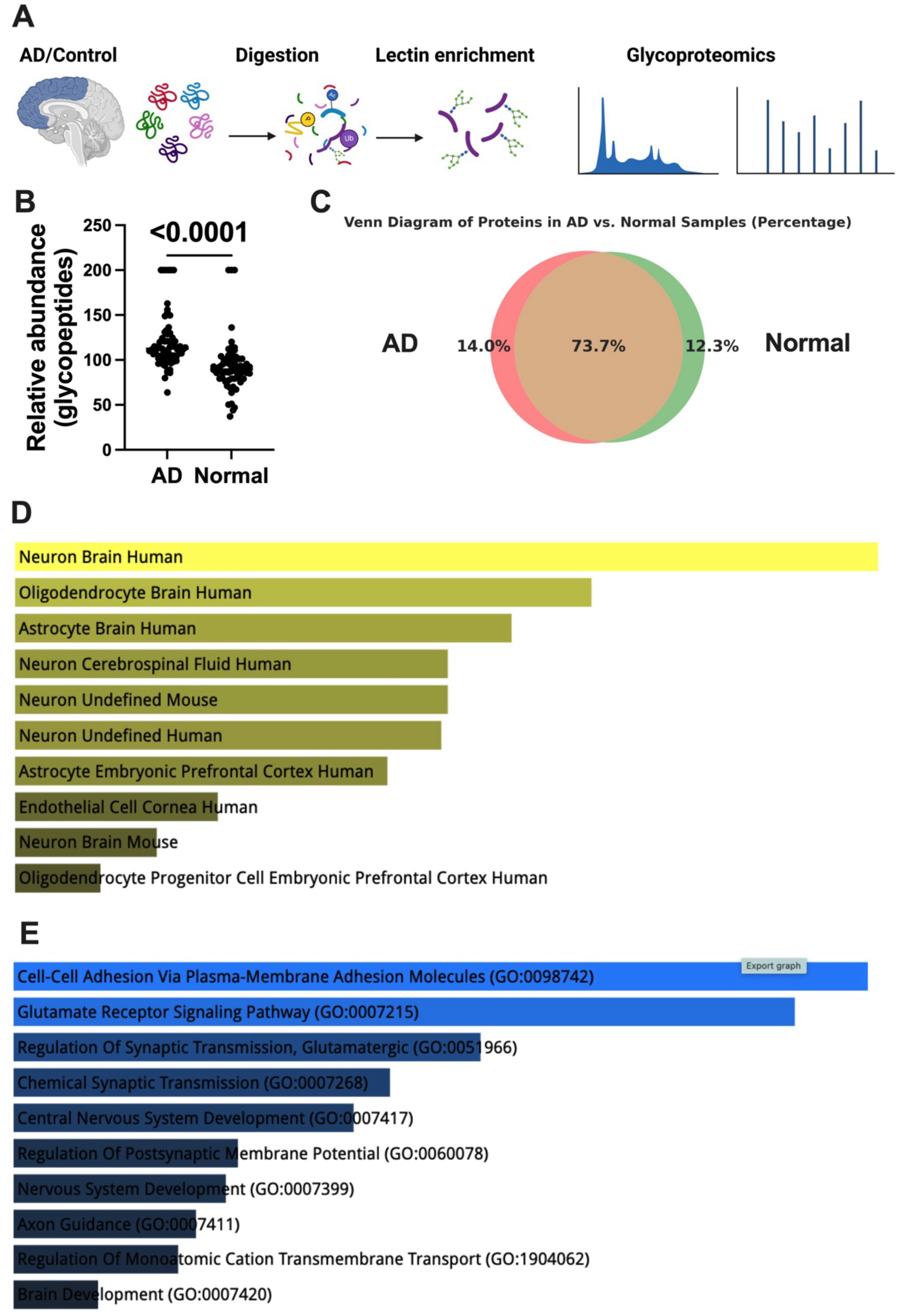
Glycoproteomic profiling reveals neuronal glycoprotein enrichment in Alzheimer’s disease mouse brains. **a**, Workflow depicting glycoproteomic analysis involving tissue homogenization, digestion, lectin-based glycopeptide enrichment, and mass spectrometry profiling from 5xFAD (AD) and WT mouse brain tissues. **b**, Quantification of total glycopeptide abundance reveals significantly elevated levels in AD samples compared to normal controls. Statistical analysis via two-tailed t-test. **c**, Venn diagram showing the overlap and distinct protein identities between AD and normal brain glycoproteomes. **d**, Cell type enrichment analysis indicates that glycoproteins upregulated in AD are predominantly expressed in neuronal and glial populations, including neurons, oligodendrocytes, and astrocytes. **e**, Gene ontology enrichment analysis reveals upregulated glycoproteins are significantly associated with neuronal functions including cell adhesion, synaptic signaling, and axon guidance.

**Extended Fig. 7.**
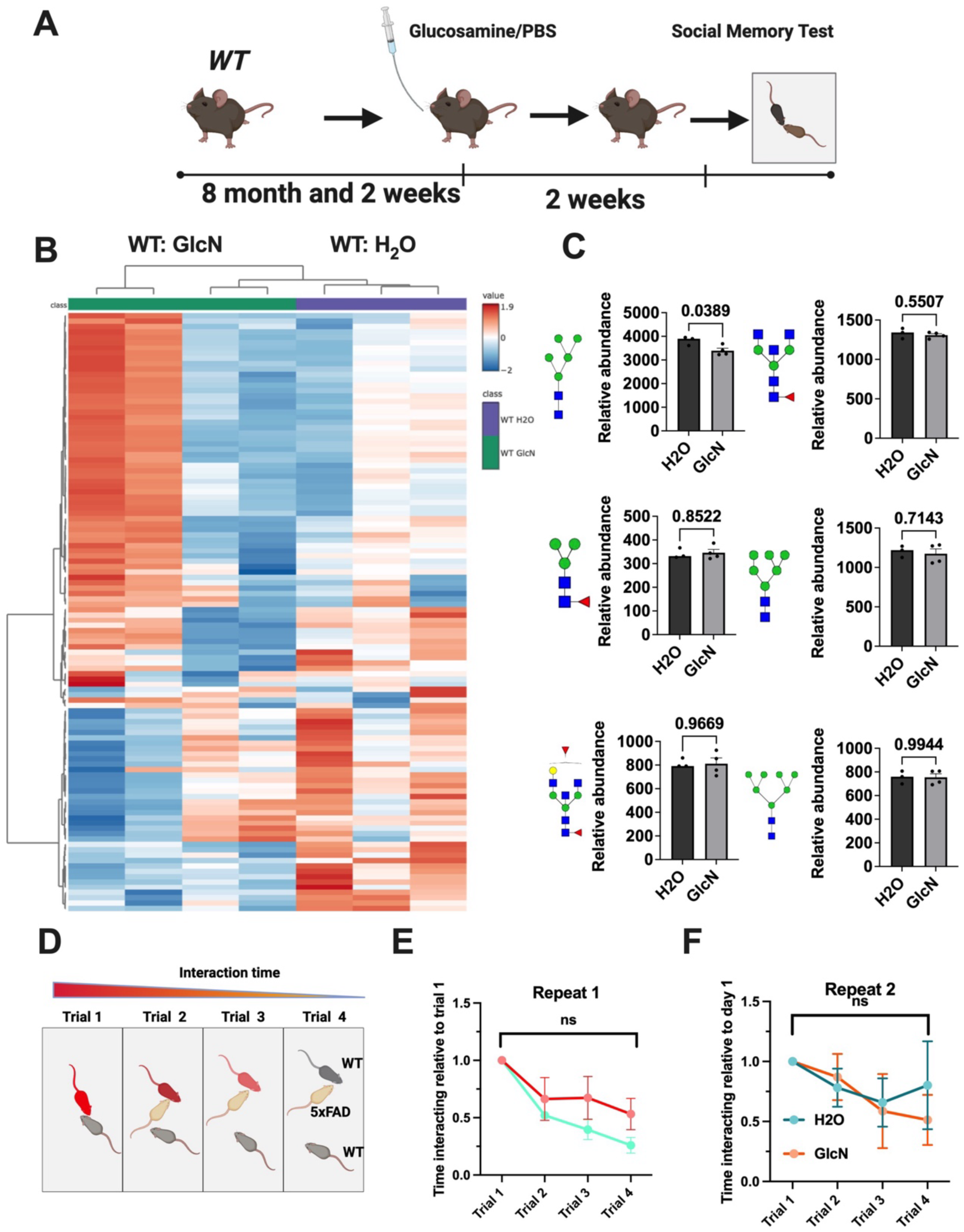
Glucosamine supplementation in wild-type mice does not induces glycan remodeling nor impairing social memory. **a**, Experimental design showing oral glucosamine (GlcN) or vehicle (H2O) supplementation in 8-month-old wild-type (WT) mice for 2 weeks, followed by social memory testing. **b**, Heatmap of glycan abundance across brain tissue from WT mice treated with GlcN or H2O, showing distinct glycan profile alterations following glucosamine supplementation. **c**, Bar graphs represent glycan species in WT GlcN versus WT H2O groups. (N=3 or 4 biological replicates per group) A subset of glycans showed significant upregulation with glucosamine treatment; statistical analysis via two-tailed t-tests; exact P-values indicated. **d**, Illustration of social memory test paradigm, with trial sequence and expected interaction dynamics. **e-f**, Social memory performance across repeated testing sessions (Repeat 1 and Repeat 2 are independent experiments) in WT mice treated with GlcN or H2O. No significant differences were observed between groups, indicating preserved cognitive function. Statistical analysis using repeated measures ANOVA.

**Extended Fig. 8.**
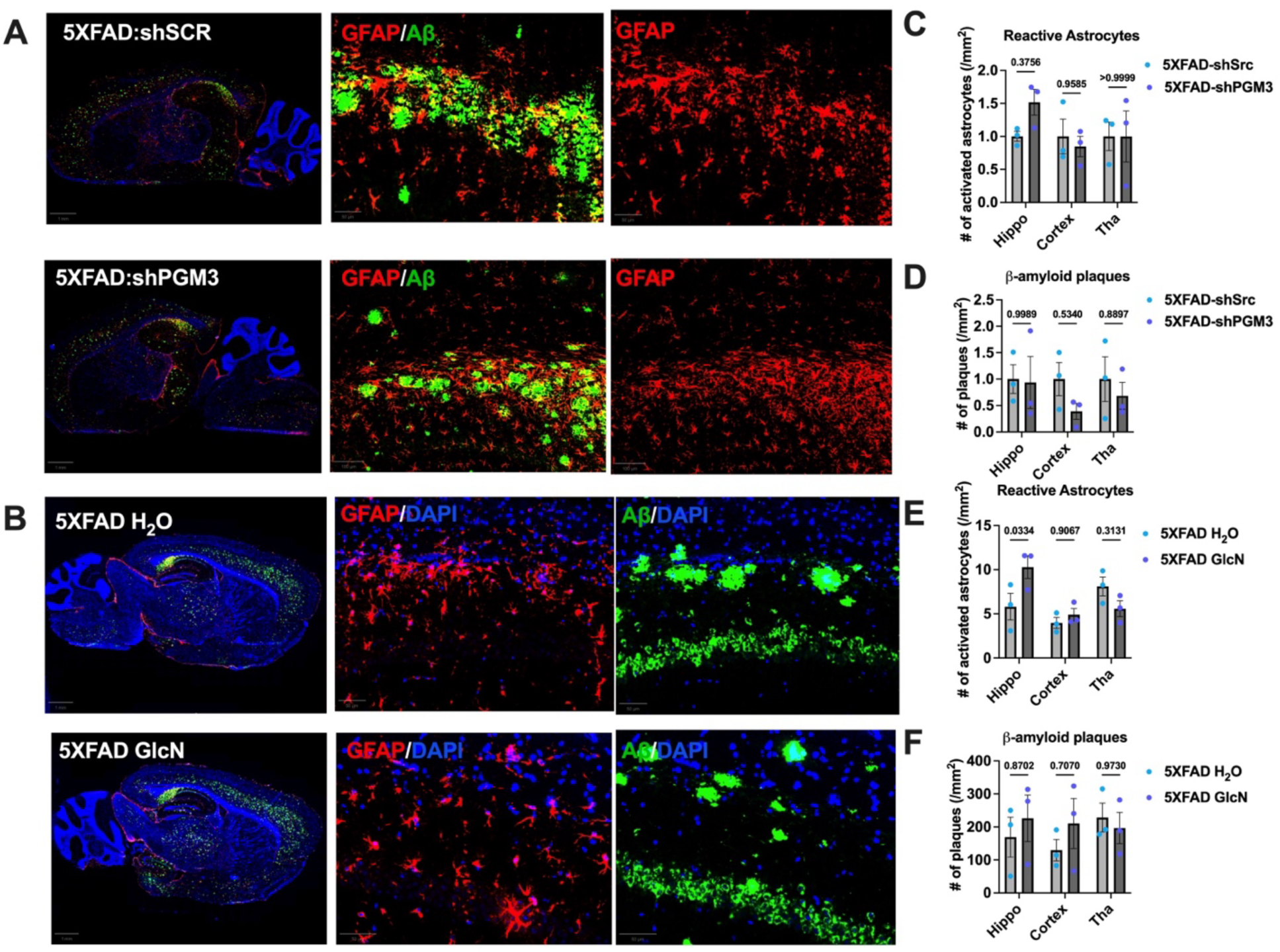
Modulating glycan biosynthesis does not affect astrocyte activation or amyloid plaque burden in 5xFAD mouse brains. **a**, Representative immunofluorescence images showing GFAP (astrocyte marker, red) and Aβ (amyloid plaques, green) staining in the hippocampus and cortex of 5xFAD mice treated with shScramble (shSCR) or shPGM3. Right panels show magnified views highlighting GFAP and Aβ signal distribution. **b**, Immunofluorescence staining of 5xFAD mice treated with glucosamine (GlcN) or water (H2O), showing GFAP (red), Aβ (green), and DAPI (blue) signals. **c-d**, Quantification of reactive astrocytes (c) and Aβ plaques (d) per mm² across hippocampus (hippo), cortex, and thalamus (tha) in 5xFAD mice treated with shPGM3 or shSCR. No significant differences were observed. Statistical analysis via two-tailed t-tests; exact P-values indicated. **e-f**, Quantification of reactive astrocytes (e) and Aβ plaques (f) in 5xFAD mice treated with GlcN or H2O. No significant differences in astrocyte activation or plaque load were detected across brain regions. Statistical analysis via two-tailed t-tests; exact P-values indicated.

## References

1 Crews, L. & Masliah, E. Molecular mechanisms of neurodegeneration in Alzheimer’s disease. Hum Mol Genet 19, R12–20 (2010).

2 Knopman, D. S. et al. Alzheimer disease. Nature Reviews Disease Primers 7, 33 (2021).

3 Golde, T. E., Schneider, L. S. & Koo, E. H. Anti-Aβ therapeutics in Alzheimer’s disease: the need for a paradigm shift. Neuron 69, 203–213 (2011).

4 Butterfield, D. A. & Halliwell, B. Oxidative stress, dysfunctional glucose metabolism and Alzheimer disease. Nature Reviews Neuroscience 20, 148–160 (2019).

5 Swerdlow, R. H. Mitochondria and Mitochondrial Cascades in Alzheimer’s Disease. J Alzheimers Dis 62, 1403–1416 (2018).

6 Di Paolo, G. & Kim, T.-W. Linking lipids to Alzheimer’s disease: cholesterol and beyond. Nature Reviews Neuroscience 12, 284–296 (2011).

7 Mosconi, L. et al. Multicenter standardized 18F-FDG PET diagnosis of mild cognitive impairment, Alzheimer’s disease, and other dementias. J Nucl Med 49, 390–398 (2008).

8 Haney, M. S. et al. APOE4/4 is linked to damaging lipid droplets in Alzheimer’s disease microglia. Nature 628, 154–161 (2024).

9 Minhas, P. S. et al. Restoring hippocampal glucose metabolism rescues cognition across Alzheimer’s disease pathologies. Science 385, eabm6131 (2024).

10 Conroy, L. R., Hawkinson, T. R., Young, L. E. A., Gentry, M. S. & Sun, R. C. Emerging roles of *N*-linked glycosylation in brain physiology and disorders. Trends in Endocrinology & Metabolism 32, 980–993 (2021).

11 Martin, P. T. Glycobiology of the synapse. Glycobiology 12, 1R–7R (2002).

12 Scott, H. & Panin, V. M. The role of protein N-glycosylation in neural transmission. Glycobiology 24, 407–417 (2014).

13 Starossom, S. C. et al. Galectin-1 deactivates classically activated microglia and protects from inflammation-induced neurodegeneration. Immunity 37, 249–263 (2012).

14 Denzel, M. S. et al. Hexosamine pathway metabolites enhance protein quality control and prolong life. Cell 156, 1167–1178 (2014).

15 Sun, R. C. et al. Brain glycogen serves as a critical glucosamine cache required for protein glycosylation. Cell Metab 33, 1404–1417.e1409 (2021).

16 Winchester, B. Lysosomal metabolism of glycoproteins. Glycobiology 15, 1R–15R (2005).

17 Freeze, H. H., Eklund, E. A., Ng, B. G. & Patterson, M. C. Neurological Aspects of Human Glycosylation Disorders. Annual Review of Neuroscience 38, 105–125 (2015).

18 Francisco, R. et al. The challenge of CDG diagnosis. Mol Genet Metab 126, 1–5 (2019).

19 Keren-Shaul, H. et al. A Unique Microglia Type Associated with Restricting Development of Alzheimer&#x2019;s Disease. Cell 169, 1276–1290.e1217 (2017).

20 de Calignon, A. et al. Propagation of tau pathology in a model of early Alzheimer’s disease. Neuron 73, 685–697 (2012).

21 Nisbet, R. M., Polanco, J.-C., Ittner, L. M. & Götz, J. Tau aggregation and its interplay with amyloid-β. Acta Neuropathologica 129, 207–220 (2015).

22 Chen, X. et al. Microglia-mediated T cell infiltration drives neurodegeneration in tauopathy. Nature 615, 668–677 (2023).

23 Shi, S. M. et al. Glycocalyx dysregulation impairs blood–brain barrier in ageing and disease. Nature (2025).

24 Han, L. et al. The role of N-glycan modification of TNFR1 in inflammatory microglia activation. Glycoconjugate Journal 32, 685–693 (2015).

25 Imbert, P. R. et al. An acquired and endogenous glycocalyx forms a bidirectional “Don’t Eat” and “Don’t Eat Me” barrier to phagocytosis. Current Biology 31, 77–89. e75 (2021).

26 Losev, Y. et al. Differential effects of putative N-glycosylation sites in human Tau on Alzheimer’s disease-related neurodegeneration. Cell Mol Life Sci 78, 2231–2245 (2021).

27 Perdivara, I. et al. Elucidation of O-Glycosylation Structures of the β-Amyloid Precursor Protein by Liquid Chromatography−Mass Spectrometry Using Electron Transfer Dissociation and Collision Induced Dissociation. J Proteome Res 8, 631–642 (2009).

28 Müller, G., Jung, C., Wied, S. & Biemer-Daub, G. Induced translocation of glycosylphosphatidylinositol-anchored proteins from lipid droplets to adiposomes in rat adipocytes. British Journal of Pharmacology 158, 749–770 (2009).

29 Hawkinson, T. R. et al. In situ spatial glycomic imaging of mouse and human Alzheimer’s disease brains. Alzheimer’s & Dementia 18, 1721–1735 (2022).

30 Ma, X. et al. AI-driven framework to map the brain metabolome in three dimensions. Nature Metabolism (2025).

31 Harrison, A. C. et al. Spatial Mapping of the Brain Metabolome Lipidome and Glycome. Nature Communications, 2023.2007.2022.550155 (2025).

32 Wang, L. et al. Spatially resolved isotope tracing reveals tissue metabolic activity. Nature Methods 19, 223–230 (2022).

33 Clarke, H. A. et al. Glycogen drives tumour initiation and progression in lung adenocarcinoma. Nature Metabolism (2025).

34 Cowie, M. R. et al. Electronic health records to facilitate clinical research. Clinical Research in Cardiology 106, 1–9 (2017).

35 Agar, N. Y. R., Yang, H. W., Carroll, R. S., Black, P. M. & Agar, J. N. Matrix Solution Fixation: Histology-Compatible Tissue Preparation for MALDI Mass Spectrometry Imaging. Analytical Chemistry 79, 7416–7423 (2007).

36 Landel, V. et al. Temporal gene profiling of the 5XFAD transgenic mouse model highlights the importance of microglial activation in Alzheimer’s disease. Molecular Neurodegeneration 9, 33 (2014).

37 Panza, F. et al. Lipid metabolism in cognitive decline and dementia. Brain research reviews 51, 275–292 (2006).

38 Mosconi, L. Brain glucose metabolism in the early and specific diagnosis of Alzheimer’s disease: FDG-PET studies in MCI and AD. European journal of nuclear medicine and molecular imaging 32, 486–510 (2005).

39 Helenius, A., Aebi & Markus. Intracellular Functions of N-Linked Glycans. Science 291, 2364–2369 (2001).

40 Seravalli, J. & Ragsdale, S. W. Pulse-Chase Studies of the Synthesis of Acetyl-CoA by Carbon Monoxide Dehydrogenase/Acetyl-CoA Synthase: EVIDENCE FOR A RANDOM MECHANISM OF METHYL AND CARBONYL ADDITION*. Journal of Biological Chemistry 283, 8384–8394 (2008).

41 Sun, R. C. et al. Noninvasive liquid diet delivery of stable isotopes into mouse models for deep metabolic network tracing. Nature Communications 8, 1646 (2017).

42 Hawkinson, T. R. & Sun, R. C. Matrix-assisted laser desorption/ionization mass spectrometry imaging of glycogen in situ. Mass Spectrometry Imaging of Small Molecules: Methods and Protocols, 215–228 (2022).

43 Du, X.-L. et al. Hyperglycemia-induced mitochondrial superoxide overproduction activates the hexosamine pathway and induces plasminogen activator inhibitor-1 expression by increasing Sp1 glycosylation. Proceedings of the National Academy of Sciences 97, 12222–12226 (2000).

44 Wang, M., Shajahan, A., Pepi, L. E., Azadi, P. & Zaia, J. Glycoproteomic Sample Processing, LC-MS, and Data Analysis Using GlycReSoft. Current Protocols 1, e84 (2021).

45 Dai, Y. et al. WebCSEA: web-based cell-type-specific enrichment analysis of genes. Nucleic acids research 50, W782–W790 (2022).

46 Zhang, Y. et al. Autosomal recessive phosphoglucomutase 3 (PGM3) mutations link glycosylation defects to atopy, immune deficiency, autoimmunity, and neurocognitive impairment. Journal of Allergy and Clinical Immunology 133, 1400–1409. e1405 (2014).

47 Rajasekaran, K. et al. Metabolic modulation of synaptic failure and thalamocortical hypersynchronization with preserved consciousness in Glut1 deficiency. Science Translational Medicine 14, eabn2956 (2022).

48 Barondes, S. H. INCORPORATION OF RADIOACTIVE GLUCOSAMINE INTO MACROMOLECULES AT NERVE ENDINGS. Journal of Neurochemistry 15, 699–706 (1968).

49 Holian, O., Dill, D. & Brunngraber, E. G. Incorporation of radioactivity of d-glucosamine-1-14C into heteropolysaccharide chains of glycoproteins in adult and developing rat brain. Archives of Biochemistry and Biophysics 142, 111–121 (1971).

50 Suzuki, K. FORMATION AND TURNOVER OF THE MAJOR BRAIN GANGLIOSIDES DURING DEVELOPMENT. Journal of Neurochemistry 14, 917–925 (1967).

51 Blömer, U. et al. Highly efficient and sustained gene transfer in adult neurons with a lentivirus vector. J Virol 71, 6641–6649 (1997).

52 Anderson, J. W., Nicolosi, R. J. & Borzelleca, J. F. Glucosamine effects in humans: a review of effects on glucose metabolism, side effects, safety considerations and efficacy. Food and Chemical Toxicology 43, 187–201 (2005).

53 Reagan-Shaw, S., Nihal, M. & Ahmad, N. Dose translation from animal to human studies revisited. The FASEB journal 22, 659–661 (2008).

54 Winslow, J. T. Mouse Social Recognition and Preference. Current Protocols in Neuroscience 22, 8.16.11–18.16.16 (2003).

55 Buglakova, E. et al. Spatial single-cell isotope tracing reveals heterogeneity of de novo fatty acid synthesis in cancer. Nature Metabolism, 1–17 (2024).

56 Alexandrov, T. Spatial metabolomics: from a niche field towards a driver of innovation. Nature Metabolism 5, 1443–1445 (2023).

57 Sun, R. C. et al. Brain glycogen serves as a critical glucosamine cache required for protein glycosylation. Cell metabolism 33, 1404–1417. e1409 (2021).

58 Piras, A., Collin, L., Grüninger, F., Graff, C. & Rönnbäck, A. Autophagic and lysosomal defects in human tauopathies: analysis of post-mortem brain from patients with familial Alzheimer disease, corticobasal degeneration and progressive supranuclear palsy. Acta Neuropathologica Communications 4, 22 (2016).

59 Nixon, R. A. & Yang, D.-S. Autophagy failure in Alzheimer’s disease—locating the primary defect. Neurobiology of Disease 43, 38–45 (2011).

60 Wang, D. et al. Glut-1 deficiency syndrome: Clinical, genetic, and therapeutic aspects. Annals of Neurology 57, 111–118 (2005).

61 Yao, J., Chen, S., Mao, Z., Cadenas, E. & Brinton, R. D. 2-Deoxy-D-Glucose Treatment Induces Ketogenesis, Sustains Mitochondrial Function, and Reduces Pathology in Female Mouse Model of Alzheimer’s Disease. PLOS ONE 6, e21788 (2011).

62 Kim, J. et al. The hexosamine biosynthesis pathway is a targetable liability in KRAS/LKB1 mutant lung cancer. Nature Metabolism 2, 1401–1412 (2020).

63 Paczynski, M. et al. The first year of amyloid immunotherapy in an academic dementia specialty practice. Alzheimer’s & Dementia 20, e086217 (2024).

64 Zeisel, S. H. Regulation of “Nutraceuticals“. Science 285, 1853–1855 (1999).

65 Shickel, B., Tighe, P. J., Bihorac, A. & Rashidi, P. Deep EHR: A Survey of Recent Advances in Deep Learning Techniques for Electronic Health Record (EHR) Analysis. IEEE Journal of Biomedical and Health Informatics 22, 1589–1604 (2018).

66 Hughes, R. & Carr, A. A randomized, double-blind, placebo-controlled trial of glucosamine sulphate as an analgesic in osteoarthritis of the knee. Rheumatology 41, 279–284 (2002).

67 Runhaar, J. et al. Subgroup analyses of the effectiveness of oral glucosamine for knee and hip osteoarthritis: a systematic review and individual patient data meta-analysis from the OA trial bank. Ann Rheum Dis 76, 1862–1869 (2017).

68 Rannou, F., Pelletier, J. P. & Martel-Pelletier, J. Efficacy and safety of topical NSAIDs in the management of osteoarthritis: Evidence from real-life setting trials and surveys. Semin Arthritis Rheum 45, S18–21 (2016).

69 Chen, Y. F. et al. Cyclooxygenase-2 selective non-steroidal anti-inflammatory drugs (etodolac, meloxicam, celecoxib, rofecoxib, etoricoxib, valdecoxib and lumiracoxib) for osteoarthritis and rheumatoid arthritis: a systematic review and economic evaluation. Health Technol Assess 12, 1–278, iii (2008).

70 Nelson, A. E., Allen, K. D., Golightly, Y. M., Goode, A. P. & Jordan, J. M. A systematic review of recommendations and guidelines for the management of osteoarthritis: The chronic osteoarthritis management initiative of the U.S. bone and joint initiative. Semin Arthritis Rheum 43, 701–712 (2014).

71 Zheng, J. et al. Association of regular glucosamine use with incident dementia: evidence from a longitudinal cohort and Mendelian randomization study. BMC Medicine 21, 114 (2023).

72 Zhou, C. et al. Habitual glucosamine use, APOE genotypes, and risk of incident cause-specific dementia in the older population. Alzheimer’s Research & Therapy 15, 152 (2023).

73 Murrey, H. E. & Hsieh-Wilson, L. C. The Chemical Neurobiology of Carbohydrates. Chemical Reviews 108, 1708–1731 (2008).

74 Leyns, C. E. G. & Holtzman, D. M. Glial contributions to neurodegeneration in tauopathies. Molecular Neurodegeneration 12, 50 (2017).

75 Kim, K. et al. Meningeal lymphatics-microglia axis regulates synaptic physiology. Cell

